# Rifampicin exposure reveals within-host *Mycobacterium tuberculosis* diversity in patients with delayed culture conversion

**DOI:** 10.1101/2020.05.22.110775

**Authors:** Charlotte Genestet, Elisabeth Hodille, Alexia Barbry, Jean-Luc Berland, Jonathan Hoffman, Emilie Westeel, Fabiola Bastian, Michel Guichardant, Samuel Venner, Gérard Lina, Christophe Ginevra, Florence Ader, Sylvain Goutelle, Oana Dumitrescu

## Abstract

*Mycobacterium tuberculosis* (Mtb) genetic micro-diversity in clinical isolates may underline mycobacterial adaptive response during the course of tuberculosis (TB) and provide insights to anti-TB treatment response and emergence of resistance. Herein we followed within-host evolution of Mtb clinical isolates in a cohort of patients with delayed Mtb culture conversion. We captured the genetic diversity of both the initial and the last Mtb isolate obtained in each patient, by focusing on minor variants detected as unfixed single nucleotide polymorphisms (SNPs). To unmask antibiotic tolerant sub-populations, we exposed the last Mtb isolate to rifampicin (RIF) prior to whole genome sequencing analysis. We were able to detect unfixed SNPs within the Mtb isolates for 9/12 patients, and non-synonymous (ns) SNPs in 6/12 patients. Furthermore, RIF exposure revealed 8 additional unfixed nsSNP unlinked to drug resistance. To better understand the dynamics of Mtb micro-diversity during the course of TB, we investigated the variant composition of a persistent Mtb clinical isolate before and after controlled stress experiments chosen to mimic the course of TB disease. A minor variant, featuring a particular mycocerosates profile, became enriched during both RIF exposure and macrophage infection. The variant was associated with drug tolerance and intracellular persistence, consistent with the pharmacological modeling predicting increased risk of treatment failure. A thorough study of such variants, not necessarily linked to canonical drug-resistance and which may be able to promote anti-TB drug tolerance, may be crucial to prevent the subsequent emergence of resistance. Taken together, the present findings support the further exploration of Mtb micro-diversity as a promising tool to detect patients at risk of poorly responding to anti-TB treatment, ultimately allowing improved and personalized TB management.

**Author summary:** Tuberculosis (TB) is caused by *Mycobacterium tuberculosis* (Mtb), bacteria that are able to persist inside the patient for many months or years, thus requiring long antibiotic treatments. Here we focused on TB patients with delayed response to treatment and we performed genetic characterization of Mtb isolates to search for sub-populations that may tolerate anti-TB drugs. We found that Mtb cultured from 9/12 patients contained different sub-populations, and *in vitro* drug exposure revealed 8 other sub-populations, none related to known drug-resistance mechanisms. Furthermore, we characterized a Mtb variant isolated from a sub-population growing in the presence of rifampicin (RIF), a major anti-TB drug. We found that this variant featured a modified lipidic envelope, and that it was able to develop in the presence of RIF and inside human macrophage cells. We performed pharmacological modelling and found that this kind of variant may be related to a poor response to treatment. In conclusion, searching for particular Mtb sub-populations may help to detect patients at risk of treatment failure and provide additional guidance for TB management.

## INTRODUCTION

Tuberculosis (TB) caused by the *Mycobacterium tuberculosis* (Mtb) complex remains one of the most prevalent and deadly infectious diseases; it was responsible for 10 million cases and 1.45 million deaths worldwide in 2018 [1]. One of the most remarkable features of Mtb infection is its chronicity, with long periods of latency, linked to the ability of the tubercle bacilli to persist in the host tissues. TB disease therefore requires a long duration of antibiotic treatment to achieve sterilization of both multiplying and dormant bacilli. Anti-TB drug resistance detection is mandatory upon TB diagnosis as it is known to hamper treatment efficacy. However, persistent infections with delayed response to treatment may be observed without any *in vitro* proven antibiotic resistance. Hypothetically, pre-existing sub-populations enclosed within Mtb clinical isolates that are antibiotic tolerant could be responsible for such persistent infections [2,3], but evidence of this is still lacking.

Since the introduction of next generation sequencing (NGS), whole genome sequencing (WGS) analysis performed on clinical Mtb isolates allowed the genetic particularities of Mtb sub-populations or variants to be revealed. Minor variants, stemmed from the initial infecting strain (hereafter referred to as the micro-diversity phenomenon), are therefore frequently present within the Mtb population before treatment onset and may also be revealed during TB treatment [4–10]. Some of these variants harbor drug resistance mutations, whilst others carry SNP in loci involved in modulation of innate immunity and in the production of Mtb cell envelop lipids [10–14]. Although the emergence of variants occurring in loci not directly related to drug-resistance is still poorly understood [15–17], this could be the key to understanding the mechanisms involved in Mtb host-adaptation possibly leading to treatment failure [18,19]. This raises the question of the evolving patterns of within-host Mtb micro-diversity in response to anti-TB treatment, and its possible link with patient outcome, beyond the canonical Mtb drug resistance phenomenon.

Delayed culture conversion (after 2 months of treatment) has been considered to be a predictor of treatment failure in TB patients [20]. We therefore focused on Mtb isolated from patients managed for drug-susceptible TB in our center between 2014 and 2016, without culture conversion after 2 months of well conducted anti-TB treatment. We captured the genetic diversity of both the initial and the last Mtb isolate obtained in each patient and we hypothesized that Mtb micro-diversity may be involved in bacilli persistence and in drug tolerance. Therefore, to unmask antibiotic tolerant sub-populations, we exposed the last Mtb isolate to rifampicin (RIF), a major anti-TB drug, prior to WGS analysis. To better understand the dynamics of Mtb micro-diversity during the course of TB, we investigated the variant composition of a persistent Mtb clinical isolate before and after controlled stress experiments chosen to mimic the course of TB disease. Importantly, a minor Mtb variant revealed after RIF exposure was found to be associated with both drug-tolerance and intra-macrophage persistence, suggesting that Mtb micro-diversity should thus be further considered to improve TB management and prevent treatment failure.

## RESULTS

### Mtb minor variants are detected during the course of TB and upon RIF exposure

From 332 susceptible TB patients managed in our center between 2014 and 2016, 16 presented positive sputum culture later than 2 months after the beginning of treatment. Four patients were lost to follow-up; 12 patients were therefore enrolled in the study (Fig 1). Ten patients presented pulmonary TB, in 2 patients TB was disseminated; all patients were smear positive. Clinical severity scores were high (Bandim TB score > 4 or Malnutrition Universal Screening Tool (MUST) > 3) in all patients. Three patients died, the other 9 patients presented no relapse at 2 years follow-up; all-but-one patient received prolonged anti-TB treatment. For each patient, genomes of both early and last Mtb isolates were analyzed with a special focus on minor variants. Moreover, to unmask RIF-tolerant sub-populations, the last isolates were exposed *in vitro* to RIF at 1x minimum inhibitory concentration (MIC) for 4 weeks prior to WGS. Among the early isolates, 3 presented 1 unfixed non-synonymous single nucleotide polymorphisms (nsSNP), while among the last isolates 3 additional unfixed nsSNP were detected, and one previously unfixed nsSNP became fixed. Interestingly, among the 8 unfixed nsSNP revealed by RIF exposure in 5 isolates none were linked to drug resistance (Table 1).

**Fig 1.**
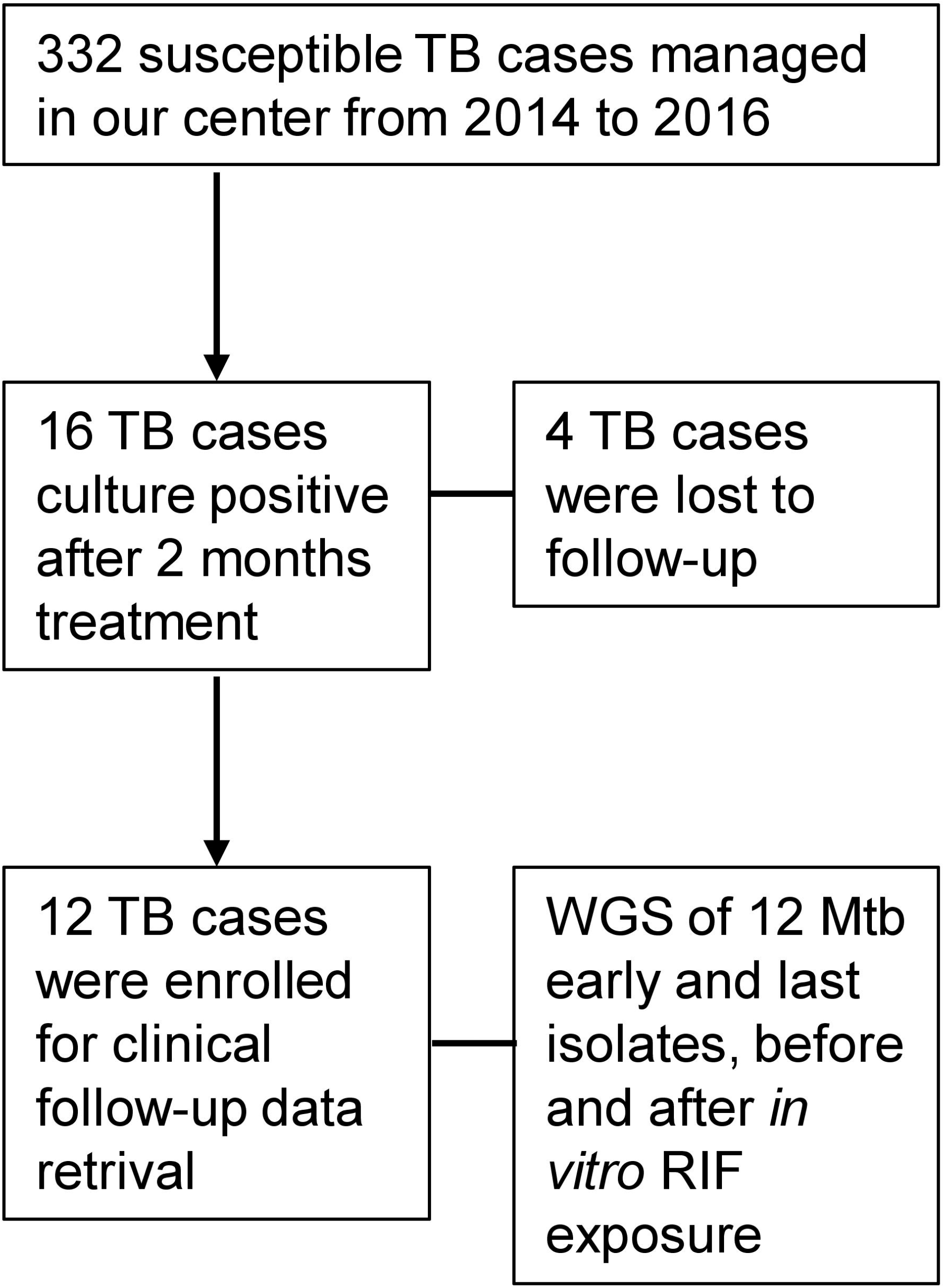
Flowchart of TB patients enrollment for WGS of early and last Mtb isolates, before and after *in vitro* rifampicin (RIF) exposure.

**Table 1.** Mtb variants identified emerging from persistent clinical isolates and after 4 weeks of exposure to 1x MIC RIF *in vitro*.

| Patient/<br>Mtb<br>lineage | Clinical<br>presentation/<br>smear | Comorbidity | Months of<br>positive<br>culture | Position | Gene | Mutation<br>type | Variant frequencies (%) |  |  | Functional<br>categories | Severity score<br>Bandim score<br>/ MUST | Outcome<br>(2 years<br>follow-up) |
| --- | --- | --- | --- | --- | --- | --- | --- | --- | --- | --- | --- | --- |
|  |  |  |  |  |  |  | Initial<br>sample | Last<br>sample | *in vitro*<br>RIF<br>exposure |  |  |  |
| P1/L2 | Disseminated<br>TB/ AFB + | Immuno-<br>suppressive<br>therapy /<br>Diabetes | M2 | 3280555 | mas Rv2940c | nsSNP | ND | ND | 44 | LM | 6 / 6 | No relapse<br>after 18<br>months anti-TB treatment |
| P2/L2 | Pulmonary<br>TB/ AFB + | None | M6 | 2892553<br>2592179 | Rv2568c<br>Rv2319c | nsSNP<br>nsSNP | ND<br>30 | ND<br>43 | 22<br>52 | CH<br>VDA | 5 / 3 | No relapse<br>after 10<br>months anti-TB treatment |
| P3/L2 | Disseminated<br>TB/ AFB + | HIV | M3 | 15876<br>2609982<br>2199585<br>899963 | pknB Rv0014c<br>cysE Rv2335<br>Rv1949c<br>cpsY Rv0806c | nsSNP<br>nsSNP<br>sSNP<br>nsSNP | ND<br>ND<br>36<br>ND | ND<br>ND<br>100<br>42 | 36<br>62<br>100<br>ND | RP<br>IMR<br>CH<br>CWCP | 7 / 4 | No relapse<br>after 12<br>months anti-TB treatment |
| P4/L3 | Pulmonary<br>TB/ AFB + | Hepatitis C | M4 | 555603<br>1803827 | Rv0465c<br>hisA Rv1603 | nsSNP<br>sSNP | ND<br>ND | 22<br>ND | ND<br>97 | IMR<br>IMR | 8 / 4 | Death after 18<br>months anti-TB treatment |
| P5/L4 | Pulmonary<br>TB/ AFB + | Immuno-<br>suppressive<br>therapy /<br>Diabetes | M2 | No<br>variant<br>identified | N/A | N/A | N/A | N/A | N/A | N/A | 7 / 4 | No relapse<br>after 9 months<br>anti-TB<br>treatment |
| P6/L4 | Pulmonary<br>TB/ AFB + | Diabetes | M2 | 328608<br>600809<br>4327302 | Rv0272c<br>hemA Rv0509<br>ethA Rv3854c | sSNP<br>sSNP<br>DEL | ND<br>ND<br>ND | 100<br>100<br>100 | 100<br>100<br>100 | CH<br>IMR<br>IMR | 5 / 2 | Death by<br>secondary<br>*Aspergillus*<br>infection |
| P7/L4 | Pulmonary<br>TB/ AFB + | Hepatitis C | M3 | 4177494 | Rv3728 | sSNP | 20 | 93 | 45 | CWCP | 9 / 6 | No relapse<br>after 9 months<br>anti-TB<br>treatment |
| P8/L4 | Pulmonary<br>TB/ AFB + | None | M2 | 3040405 | miaA Rv2727c | nsSNP | ND | ND | 20 | IMR | 6 / 6 | Death before<br>treatment<br>completion |
| P9/L4 | Pulmonary<br>TB/ AFB + | None | M2 | 115217<br>966395 | nrp Rv0101<br>moaA Rv0869c | sSNP<br>nsSNP | 25<br>22 | ND<br>ND | ND<br>ND | LM<br>IMR | 2 / 5 | No relapse<br>after 9 months<br>anti-TB<br>treatment |
| P10/L4 | Pulmonary<br>TB/ AFB + | None | M2 | 853811<br>3771284<br>1150503 | phoR Rv0758<br>relK Rv3358<br>kdpD Rv1028c | sSNP<br>sSNP<br>nsSNP | ND<br>ND<br>ND | 35<br>29<br>56 | 20<br>18<br>ND | RP<br>VDA<br>RP | 5 / 4 | No relapse<br>after 6 months<br>anti-TB<br>treatment |
| P11/L4 | Pulmonary<br>TB/ AFB + | None | M2 | 183015<br>379185 | fadE2 Rv0154c<br>Rv0311 | sSNP<br>nsSNP | 15<br>17 | 100<br>100 | 100<br>100 | LM<br>CH | 7 / 4 | No relapse<br>after 9 months<br>anti-TB<br>treatment |
| P12/L4 | Pulmonary<br>TB/ AFB + | None | M3 | 451849<br>1689565<br>3513662 | Rv0374c<br>Rv1498c<br>nuoD Rv3148 | nsSNP<br>nsSNP<br>nsSNP | ND<br>ND<br>ND | ND<br>ND<br>ND | 36<br>21<br>20 | IMR<br>IMR<br>IMR | 5 / 4 | No relapse<br>after 9 months<br>anti-TB<br>treatment |

P: Patient; L: Mtb lineage; AFB: Acid Fast Bacilli; RIF: Rifampicin; N/A: Not Applicable; ND: Not detected = Below Limit of Detection;
Conserved hypotheticals: CH; Cell wall and cell processes: CWCP; Intermediary metabolism and respiration: IMR; Lipid metabolism: LM;
Regulatory proteins: RP; Virulence, detoxification, adaptation: VDA.

### A minor variant of Mtb clinical isolate is selected in response to both rifampicin exposure and macrophage infection

We focused on the P1 strain (Table 1) in which RIF exposure revealed a single non-fixed variant (C 3280555 G), leading to a nsSNP in a locus with known function involved in lipid metabolism. To decipher the dynamics of the variant composition of this clinical isolate we first performed WGS on 20 colonies. Apart from the predominant clone, 4 variants were identified and their presence in the initial isolate was confirmed by targeted NGS, at frequencies between 0.5% and 9% (Fig 2A). Six loci differed among the variants, with a maximum pairwise distance of 4 SNPs. On the one hand, the initial isolate was exposed for 4 weeks to RIF, *in vitro*, at 1xMIC (Fig 2B) and on the other hand it was submitted to 7 days of macrophage infection (Fig 2C). Interestingly, NGS analysis found that the C 3280555 G variant was the most enriched during both RIF exposure (from 1.5% to 45%) and macrophage infection (from 1.5% to 74%), while the relative frequency of the other 3 variants varied within a range of 0.05 to 10% (Fig 2).

**Fig 2.**
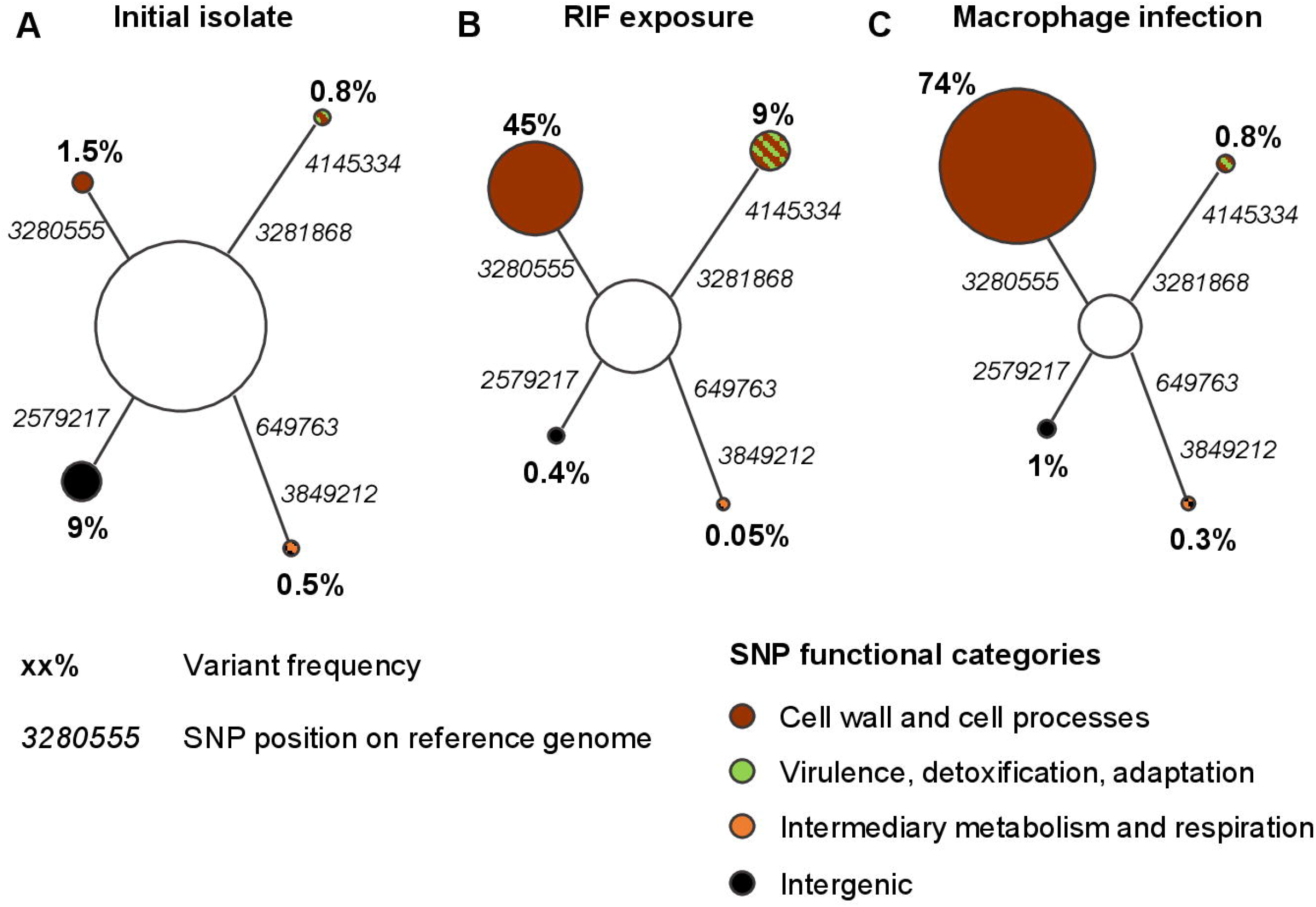
A minor variant of Mtb clinical isolate is selected in response to both rifampicin (RIF) exposure and macrophage infection. Minimum spanning trees (MST) of (A) the initial clinical isolate, (B) after 4 weeks of *in vitro* RIF exposure at 1x minimum inhibitory concentration (MIC) or (C) after 7 days of macrophage infection at a multiplicity of infection (MOI) of 10:1 (bacteria:cells) assessed by targeted NGS with coverage ranging from 10 000 to 36 000x. MST are pooled results of two independent experiments.

We further studied the dynamics of the diversity of the clinical isolate by focusing on its two main composing variants that we discriminated, according to WGS data, by the nucleotide 3280555 (i.e. G for the minor variant and C for the major variant). Two cloned variants stemmed from the initial clinical isolates were compared: the initially majority variant (IMV), carrying the C nucleotide at position 3280555, and the variant carrying the fixed C 3280555 G polymorphism. When appropriate, experiments were also performed on a 50:50 mixture of these two variants.

### The C 3280555 G harboring variant overexpresses mycocerosates and more particularly the tetramethyl-branched components of mycocerosates

The SNP C 3280555 G results in an amino-acid change, A721P, of mycocerosic acid synthase (Mas, Rv2940c). It is located in the Mas acyltransferase domain, which is involved in multimethylated mycocerosates synthesis by iterative condensation of methyl-malonyl-CoA units [21,22]. Mycocerosates are components of the phthiocerol dimycocerosate (PDIM) and of phenolic glycolipids (PGL) of Mtb, both strongly involved in host-pathogen interaction [23–25]. Because the mycocerosate components are expected to be altered in the C 3280555 G variant, the lipid profile of this variant and of the IMV were explored, focusing on the production of the 4 main components of mycocerosates (Fig 3A): the two trimethyl-branched, C27 and C29, and the two tetramethyl-branched, C30 and C32 [26,27]. When compared to the IMV, the C 3280555 G variant profile showed significantly lower trimethyl-branched C27 component (11.4 ± 2.2%) than tetramethyl-branched C30 component (80.1 ± 2.4%; Fig 3B), consistent with a higher affinity of Mas enzyme for methyl-malonyl-CoA that promotes tretramethyl mycocerosate synthesis in this variant. Moreover, the relative proportion of mycocerosates produced by the C 3280555 G variant was approximately twice as high as that produced by the IMV (Fig 3C). Given this lipid profile, the variant carrying the C 3280555 G polymorphism is hereafter referred to as the 4MBE (<u>tetra</u>-<u>m</u>ethyl <u>b</u>ranched <u>e</u>nriched) variant.

**Fig 3.**
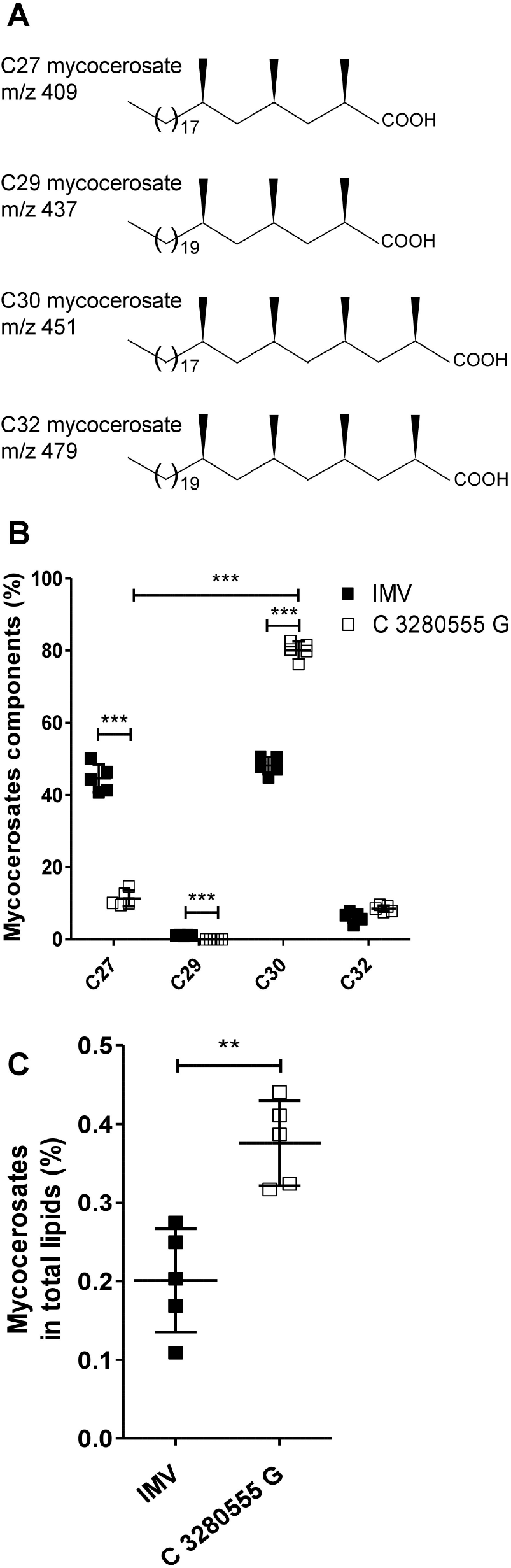
The 4MBE variant overexpresses mycocerosates and more particularly the tetramethyl-branched components of mycocerosates. (A) Molecular structures of C27, C29, C30 and C32 mycocerosate components from Mtb. (B-C) The lipid profile of the initially majority variant (IMV, black) and the C 3280555 G variant (white) was assessed. (B) Proportion of mycocerosate components regarding total mycocerosates were explored by gas chromatography-mass spectrometry (GC-MS) and obtained from ion chromatograms showing the fragment ions at m/z 409, 437, 451 and 479 representing mycocerosate components C27, C29, C30 and C32, respectively. (C) Proportion of mycocerosates in total lipids were obtained from ion chromatograms showing all fatty acids detected by GC-MS. Values for each condition are the mean ± standard deviation (SD) of five independent experiments. Means were compared using One-Way ANOVA followed by Bonferroni correction (B) or using unpaired t-test (C).

### The 4MBE variant is tolerant to RIF alone and in combination with isoniazid

As the 4MBE variant was revealed upon RIF exposure its susceptibility to this drug was explored. Although there was no difference regarding the MIC values (0.12 mg/L for both 4MBE and IMV), the tolerance of the 4MBE variant to sub-MIC doses of RIF was investigated. The kinetics of the bacterial growth, explored by the time to positivity (TTP) approach, revealed similar growth profiles in the absence of RIF (from 6.7 ± 0.3 days to 7.2 ± 0.3 days), showing the same fitness of these variants in the absence of antibiotic pressure. Conversely, a significantly shorter TTP for the 4MBE variant (7.1 ± 0.4 days) compared to the IMV (15.6 ± 1.3 days) was observed upon antibiotic treatment with 1xMIC of RIF (Fig 4A), suggesting a better ability of the 4MBE variant to persist upon RIF exposure.

**Fig 4.**
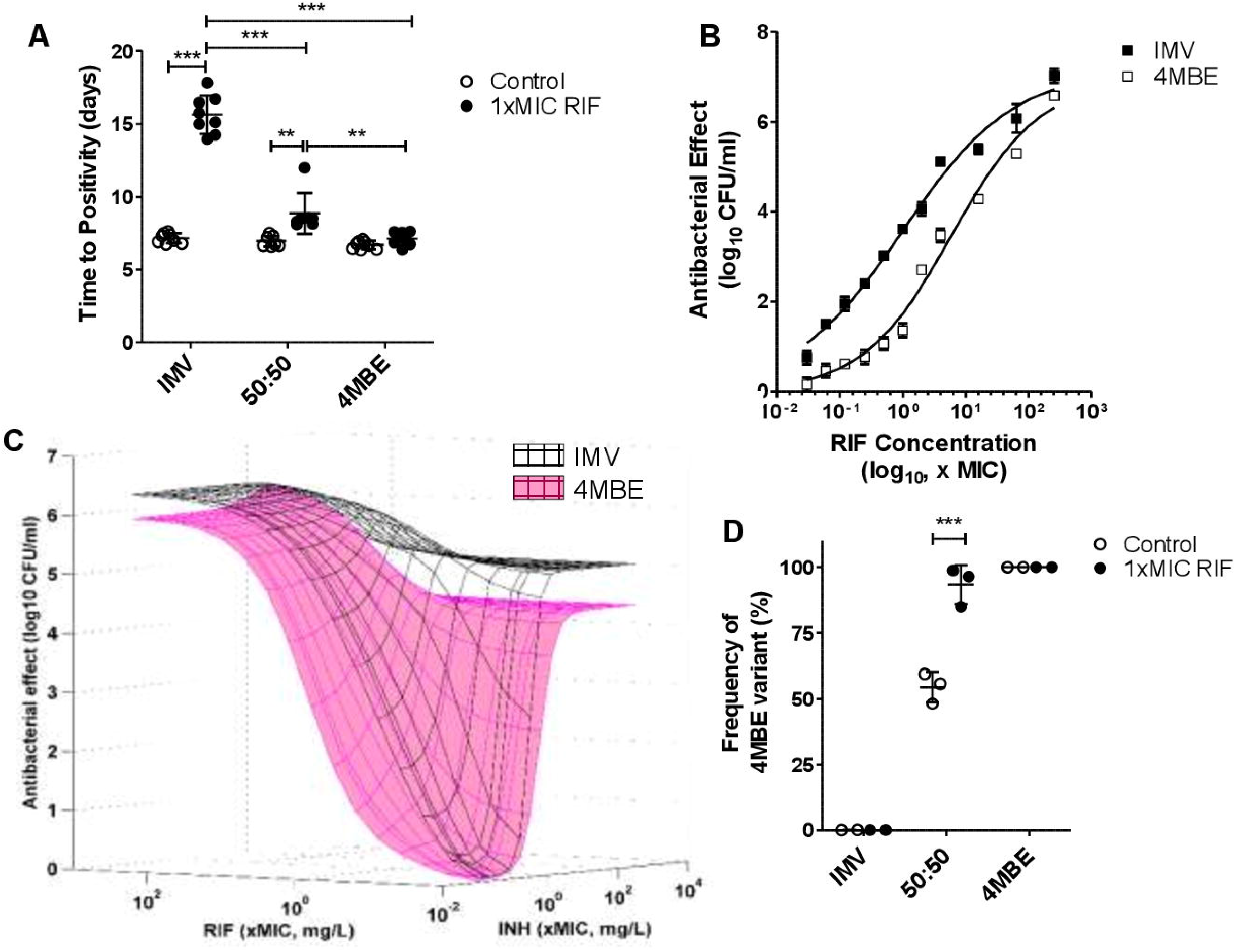
The 4MBE variant is tolerant to rifampicin (RIF) alone and in combination with isoniazid (INH). (A) Mycobacterial growth of the IMV, 4MBE variant, and a 50:50 mixture in absence (white; Ctrl) or presence of RIF (black; 1x minimum inhibitory concentration (MIC) RIF) is expressed in time to positivity (TTP) in the BACTEC system. Values for each condition are the mean ± standard deviation (SD) of seven or eight independent experiments. Means were compared using one-way ANOVA followed by Bonferroni correction. (B) Time-kill data were determined after 7 days of RIF exposure, at concentrations ranging from 1/32 to 256xMIC, for the IMV (black) and the 4MBE variant (white). Fit of the Hill pharmacodynamic model to rifampicin (RIF) time-kill data, the solid lines are the best fit lines. The y-axis represents the difference in log_10_ CFU/mL relative to control (C = 0) from a duplicate experiment. (C) Superposition of response surface models of the combined antibacterial effect of INH and RIF, from 1/32 to 256xMIC, against the IMV (black) and the 4MBE variant (pink). A total of 142 measures of the effect of RIF and INH alone or in combination were performed, and results were analyzed using the Minto response surface model. The surfaces represent the model-based predicted effect, representing the antibacterial effect measured after 7 days of exposure. (D) After 4 weeks of incubation without (white; Ctrl) or with RIF (black; 1xMIC RIF), the change in the 4MBE variant frequency on plated bacteria was assessed by droplet digital PCR (ddPCR) as described in the Methods section. Values for each condition are the mean ± standard deviation (SD) of two or three independent experiments. Means were compared using the Student’s t-test.

To further explore RIF tolerance of the 4MBE variant, concentration-effect experiments were performed, and a pharmacological efficacy prediction model was then applied to the results obtained. While there were no significant differences for the maximal effect of RIF, there was a decrease in RIF susceptibility for the 4MBE variant at sub-MIC doses (Fig 4B). The 4MBE variant exhibited higher EC50 value (5.9, 95% CI [3.4; 10.4]) than the IMV (1.0, [0.6; 1.6]), suggesting a lower potency of this antibiotic on 4MBE variant (S1 Table). Overall, these results indicate that the 4MBE variant is more tolerant to sub-MIC doses of RIF compared to the IMV, despite the absence of canonical RIF resistance.

As RIF is part of a complex regimen during TB treatment, the antibacterial effect of the combination of both first-line anti-TB drugs (isoniazid [INH] and RIF) was investigated on the IMV and the 4MBE variant (Fig 4C). The Minto model described the combined antibacterial effect very well, R^2^ values ≥ 0.90 and low bias and imprecision values (S2 Table). For RIF an almost 6-fold higher EC50 value was observed for the 4MBE variant compared to the IMV. For INH, the 4MBE variant exhibited a slightly higher EC50 value (0.91 [0.80; 1.03]) than the IMV (0.53 [0.48–0.58]), and exhibited also a lower Hill coefficient value (2.07 [1.74; 2.41]) than the IMV (3.54 [2.64, 4.43]), suggesting also a lower potency of INH on the 4MBE variant. More importantly, the interaction parameter βU50 was significantly greater than 0 for the IMV (1.43 [0.94; 1.91]), which means that the combined effect of INH and RIF was synergistic. Conversely, it was not significantly different from 0 for the 4MBE variant (0.68 [-0.38; 1.74]), indicating that the combined effect was only additive. Taken together, these results indicate that the 4MBE variant have increased tolerance to RIF and INH, alone but also in combination, possibly favoring treatment failure.

To explore a possible selective advantage of the 4MBE variant upon treatment, the dynamics of variant assemblies upon RIF exposure was monitored. Mycobacteria in a IMV:4MBE 50:50 mixture were recovered after 4 weeks of growth in MGIT culture tubes, with or without 1xMIC of RIF, to extract DNA after plating and assess the frequency of the 4MBE variant by ddPCR (Fig 4D). Without antibiotic pressure, the distribution of the variants in the 50:50 mixture was substantially stable (54.4 ± 5.7 %) after 4 weeks of culture. Conversely, after RIF exposure, the 4MBE variant became predominant (93.4 ± 7.3%), consistent with the previous NGS experiments (Fig 2B).

### Improved Mtb intra-macrophagic survival of the 4MBE variant

As the 4MBE variant was also revealed upon macrophage infection (Fig 2C), the abilities of intra-macrophage survival of the IMV, the 4MBE variant and their 50:50 mixture were explored at 6 h and 96 h post-infection (hpi).

At 6 hpi, the ability of the 4MBE variant to invade macrophages was significantly better (28.1×10^4^ ± 9.0×10^4^ CFU/mL) than that of the IMV (4.7×10^4^ ± 2.4×10^4^ CFU/mL, Fig 5A). Moreover, the intracellular growth ratio between 96 hpi and 6 hpi was significantly better for the 4MBE variant (5.0 ± 0.9-fold) than for the IMV (1.4 ± 0.4-fold), indicating a better intra-macrophagic survival and/or multiplication (Fig 5B).

**Fig 5.**
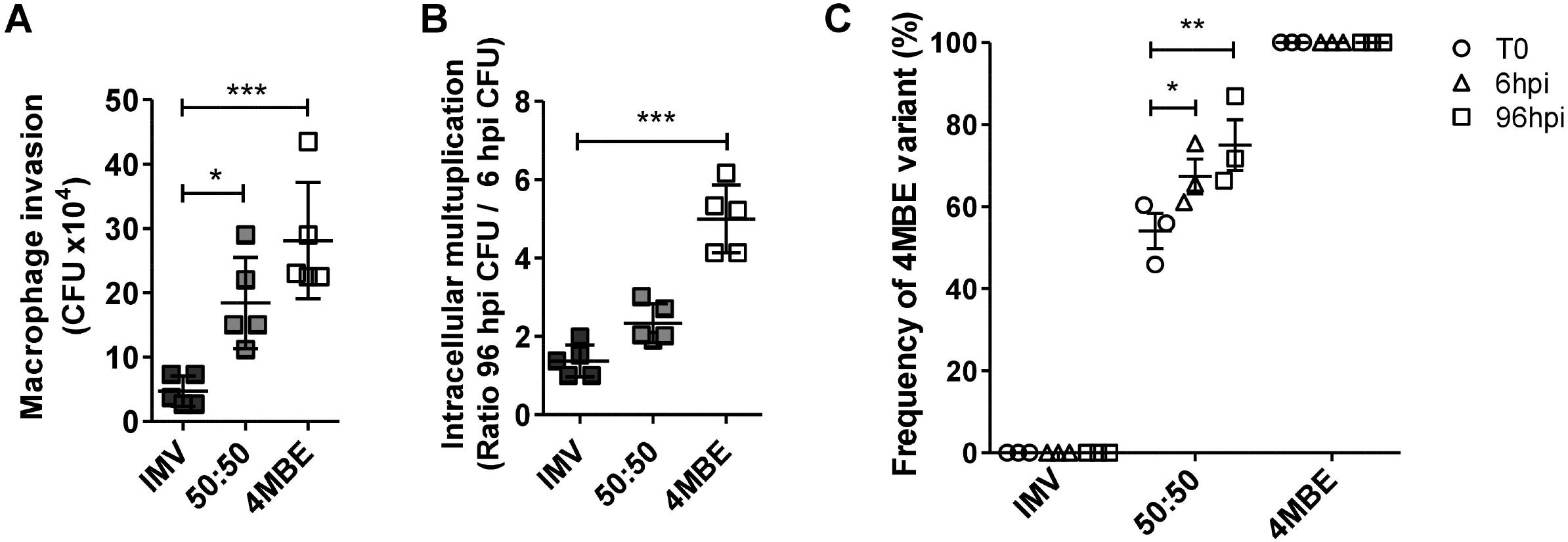
Improved Mtb intra-macrophagic survival of the 4MBE variant. Macrophages were infected by the IMV (black), the 4MBE variant (white) or a half-half mixture (50:50, grey) at a multiplicity of infection (MOI) of 10:1. (A-B) At 6 and 96 hours post-infection (hpi), macrophages were lysed for intracellular CFU counting to study (A) macrophage invasion thanks to intracellular CFU counting at 6 hpi and (B) intracellular multiplication thanks to intracellular CFU ratio between 6 hpi and 96 hpi. (C) The change in the 4MBE variant frequency on plated bacteria was assessed by ddPCR as described in the Methods section. Values for each condition are the mean ± SD of three to five independent experiments. Means were compared using One-Way ANOVA followed by Bonferroni correction.

To further monitor the relative proportion between the variants upon host-pathogen interaction, the DNA of plated bacteria at 6 hpi and 96 hpi were recovered to assess the variant frequency by ddPCR (Fig 5C). An enrichment of the 4MBE variant, up to 75.0 ±10.7% at 96 hpi, was observed, consistent with the previous NGS experiments (Fig 2C).

### Improved intra-macrophagic survival of the 4MBE variant is due to phagolysosome avoidance and inhibition of autolysosome formation

Mtb intramacrophagic behavior of the IMV and the 4MBE variant was explored by confocal microscopy analysis at 24 hpi (Fig 6). The improved Mtb intra-macrophagic survival upon infection with the 4MBE variant compared to the IMV was confirmed by a significantly greater number of both infected cells and bacterial burden per cell (S1 Fig).

**Fig 6.**
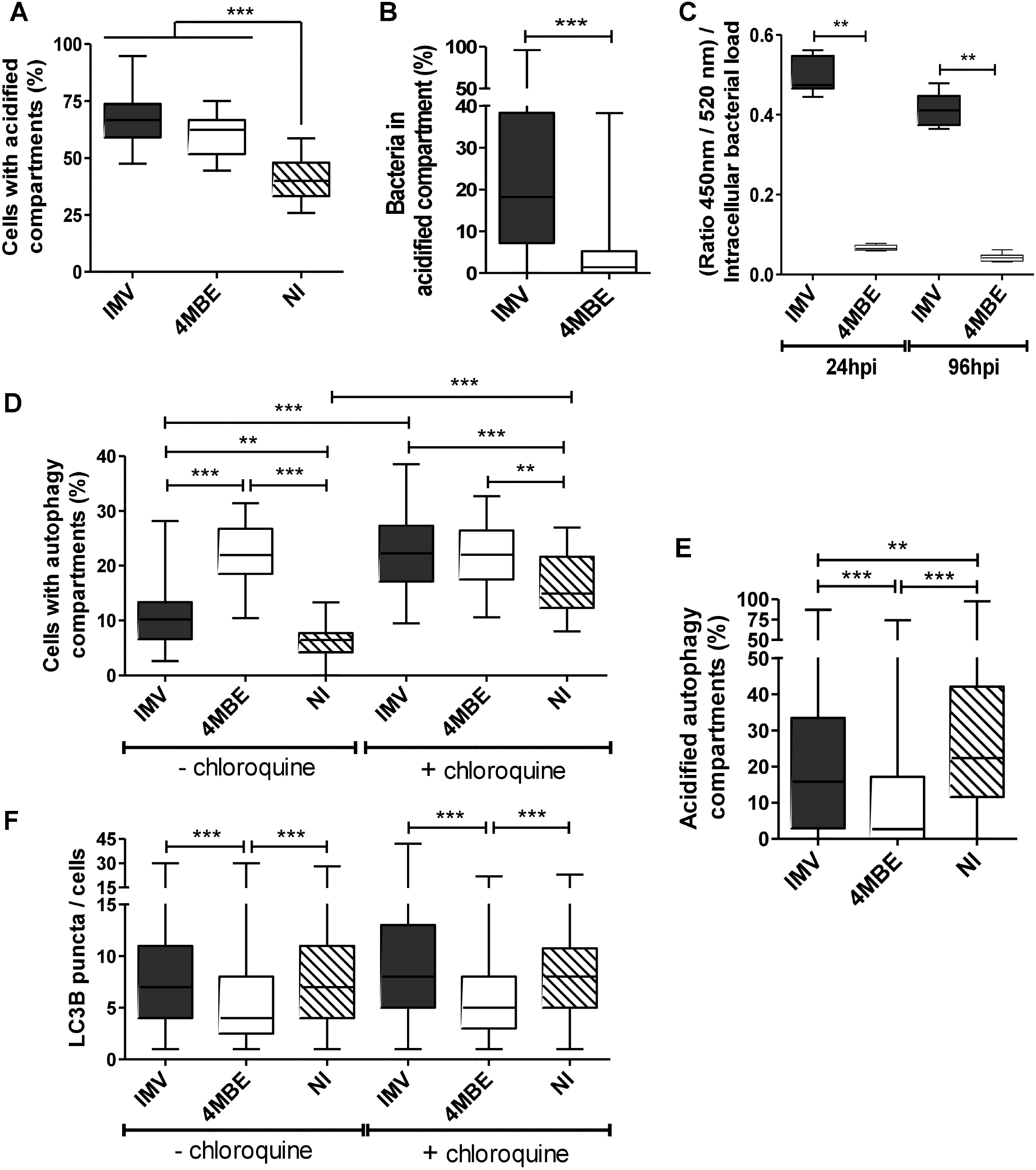
Improved intra-macrophagic survival of the 4MBE variant is due to phagolysosome avoidance and inhibition of autolysosome formation. Macrophages were infected by the IMV (black) or the 4MBE variant (white) at a MOI of 10:1 or not infected (NI; hatched) as a control. (A-B and D-F) At 24 hpi, cells were stained with LysoTracker Red DND-99, anti-Mtb coupled FITC, and anti-LC3B coupled Dy-Light 650 antibodies. Cells with acidified (A) or autophagy compartments (D) were determined across at least 10 confocal images, containing between 30 and 80 cells per image, per replicate. Proportion of bacteria in acidified compartments (B), proportion of acidified autophagy compartments (E), and LC3B puncta per cells (F) were determined across at least 40 cells per replicate. (D and F) Three hours before staining, cells were incubated with chloroquine, then macrophages were stained and analyzed as previously described. (A-B and D-F) Values for each condition are the median values [interquartile range, IQR] of at least three independent experiments. Statistical significance was determined using Kruskal-Wallis analysis, using Dunn’s Multiple Comparison Test or Mann Whitney test, where appropriate. (C) Infected macrophages were loaded with CCF4-AM at 24 and 96 hpi to analyze 450nm/520nm ratio normalized to bacterial load. Values for each condition are the mean ± standard deviation (SD) of at least three independent experiments. Means were compared using Mann Whitney test.

To explore the mechanisms of intra-macrophage survival of the 4MBE variant, phagolysosome activation and avoidance were first explored (Fig 6A-C). Although no difference was observed in the number of cells with acidified compartments (Fig 6A), there were significantly more bacteria in acidified compartments upon infection by the IMV (18.2% [7.1-38.3]) compared to the 4MBE variant (1.4% [0.03-5.2], Fig 6B). To determine whether this decreased colocalization of Mtb with acidified compartments was due to Mtb translocation from the phagolysosome to the cytosol or to an inhibition of phagolysosomal fusion, infected macrophages were loaded with CCF4-AM, a fluorescence resonance energy transfer (FRET)-based assay. After normalization on the intracellular bacterial load, the blue/green signal was almost 10 times lower upon infection by the 4MBE variant than the IMV at 96 hpi (Fig 6C). This supports the notion that the 4MBE variant avoids phagolysosomal fusion rather than translocating to the cytosol.

The autophagy activation and outcome were then explored. Although the proportion of cells with autophagy compartments was approximately two-fold upon infection by the 4MBE variant compared to the IMV, chloroquine treatment abolished the differences between the infecting variants. Moreover, the presence or absence of chloroquine pretreatment did not modify the proportion of cells with autophagy compartments upon infection by the 4MBE variant, suggesting an inhibition of the autolysosome formation by this variant (Fig 6D). This was confirmed by the significant decrease in the proportion of acidified autophagy compartments upon infection by the 4MBE variant compared to the IMV (Fig 6E). Furthermore, there was a decrease in the number of LC3B puncta per cells upon infection by the 4MBE variant, of approximately 1.5-fold, compared to the IMV (Fig 6F), indicating a weaker autophagy activation upon infection by the 4MBE variant.

Taken together, these results support that the better intracellular survival of the 4MBE variant during macrophage infection is driven by the increase of phagolysosome avoidance and the decrease of autophagy activation and outcome.

### Macrophage apoptosis and inflammatory response are exacerbated upon infection by the 4MBE variant

Macrophage transcriptome analysis found that 342 genes were differentially expressed between macrophages infected by the IMV and those infected by the 4MBE variant. A total of 228 differentially expressed genes were selected to build a heatmap, showing clustering according to biological replicates (Fig 7). The differentially expressed genes were classified into nine functional categories involved in Mtb macrophage response. Consistent with the macrophage infection experiments, actin cytoskeleton, autophagy, and phagolysosome pathways were found to be dysregulated upon infection by the 4MBE variant compared to the IMV. Moreover, there was also a dysregulation of inflammatory response, apoptosis and cell cycle, mitochondrial metabolism, lipid metabolism, and T cell signaling pathways.

**Fig 7.**
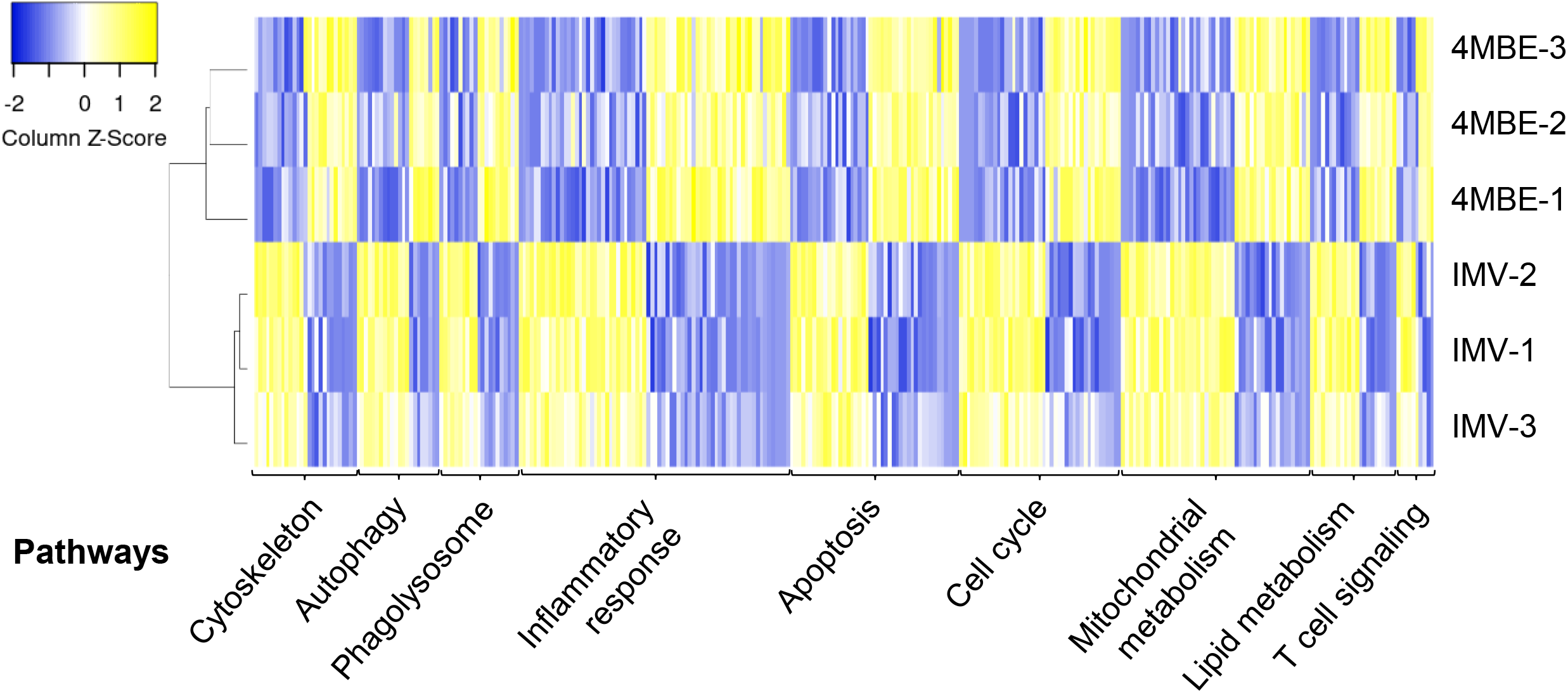
Dysregulation of macrophage gene expression upon infection by the 4MBE variant. Macrophages were infected at a MOI of 10:1 by the IMV or the 4MBE variant. At 24 hpi, total RNA were extracted to study the changes in global transcriptome of host cells. Heatmap of gene expression (log_2_ fold change) for 228 differentially expressed genes between IMV or 4MBE variant-infected macrophages at 24 hpi. Rows are independent experiments and columns are genes. Differentially expressed genes were classified in nine functional categories. Clustering was based on Pearson’s correlation. The two experimental conditions clustered with their biological replicates.

In order to explore cell death mechanisms, lytic cell death (Fig 8A) and apoptosis (Fig 8B) of macrophages were assessed upon infection by the IMV and the 4MBE variant at 24 and 96 hpi, as apoptosis is generally regarded as a protective response, whereas lytic cell death is thought to favor inflammation and disease progression [28–30]. At 96 hpi, the lytic cell death upon 4MBE variant infection (64.9 ± 3.5%) was significantly greater than that obtained upon IMV infection (Fig 8A). Furthermore, apoptosis at 96hpi was also significantly higher upon infection by the 4MBE variant (15.1% [11.1-19.2]) compared to the IMV (7.9% [4.7-12.0], Fig 8B).

**Fig 8.**
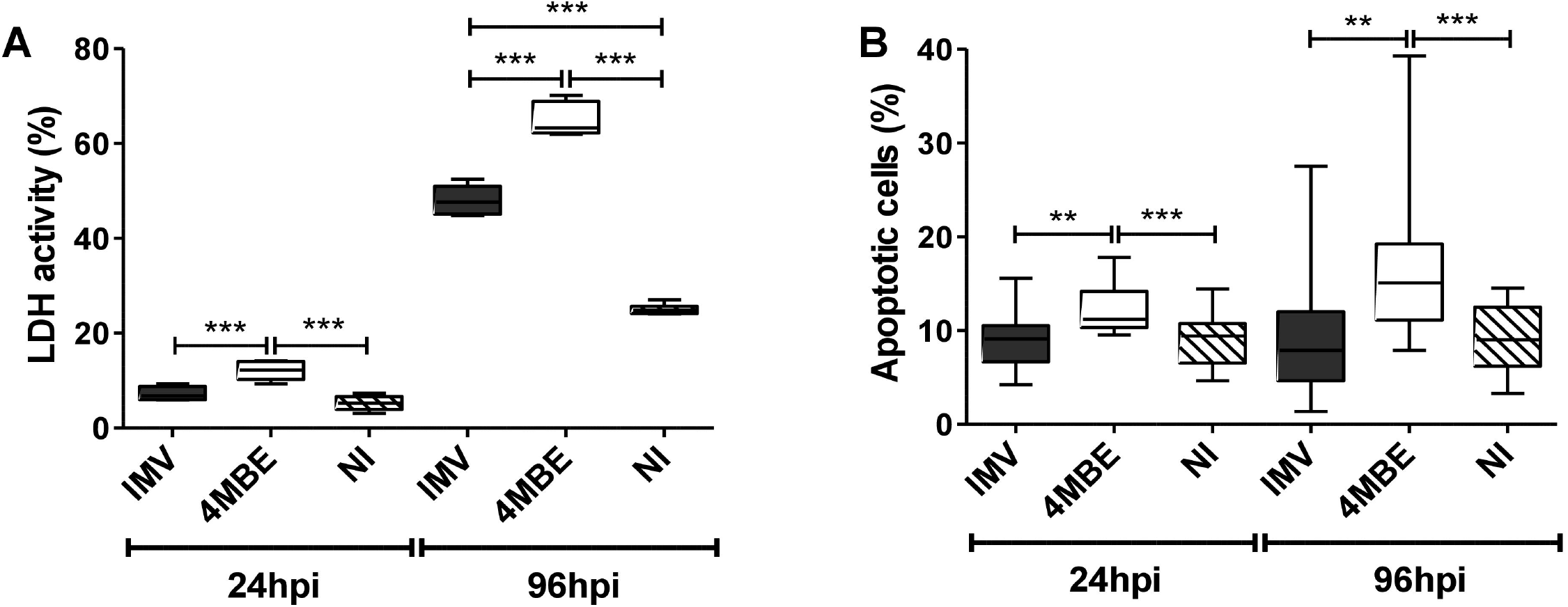
Exacerbated macrophage cell death upon infection by the 4MBE variant. Macrophages were infected by the IMV (black) or the 4MBE variant (white) at a MOI of 10:1 or not infected (NI; hatched) as a control. (A) Lytic cell death, at 24 hpi and 96 hpi, was evaluated by LDH release in cell culture supernatant from at least 3 independent experiments analyzed in duplicate. Means were compared using One-Way ANOVA followed by Bonferroni correction. (B) Apoptosis was measured by Annexin V staining and analysis performed by microscopy, at 24 hpi and 96 hpi. Apoptosis was determined across at least 5 images, containing between 20 and 50 cells per image, per replicate, in at least three independent experiments. Statistical significance was determined using Kruskal-Wallis analysis, using Dunn’s Multiple Comparison Test.

Cytokine (TNF-α, IL-6, IL-1β, IFN-γ), chemokine (RANTES (CCL5), and interferon gamma-induced protein 10 (IP-10) production were then measured at 6 hpi (S2 Fig), 24 hpi (Fig 9), and 96 hpi (S3 Fig) to explore macrophage inflammatory response. At 24hpi, there was a significant increase in the inflammatory response of macrophages upon infection by the 4MBE variant compared to the IMV (Fig 8). Upon infection by the 4MBE variant there was a significant increase in TNF-α production by macrophages from 6 to 96 hpi (Fig 9A, S2A-S3A Figs) and overall a sustained inflammatory response at 96 hpi (Fig S3). Finally, chemokine signals were significantly increased upon infection by the 4MBE variant (Fig 9E-F). Overall, the macrophage pro-inflammatory response was stronger (in terms of intensity and duration) upon infection by the 4MBE variant than by the IMV.

**Fig 9.**
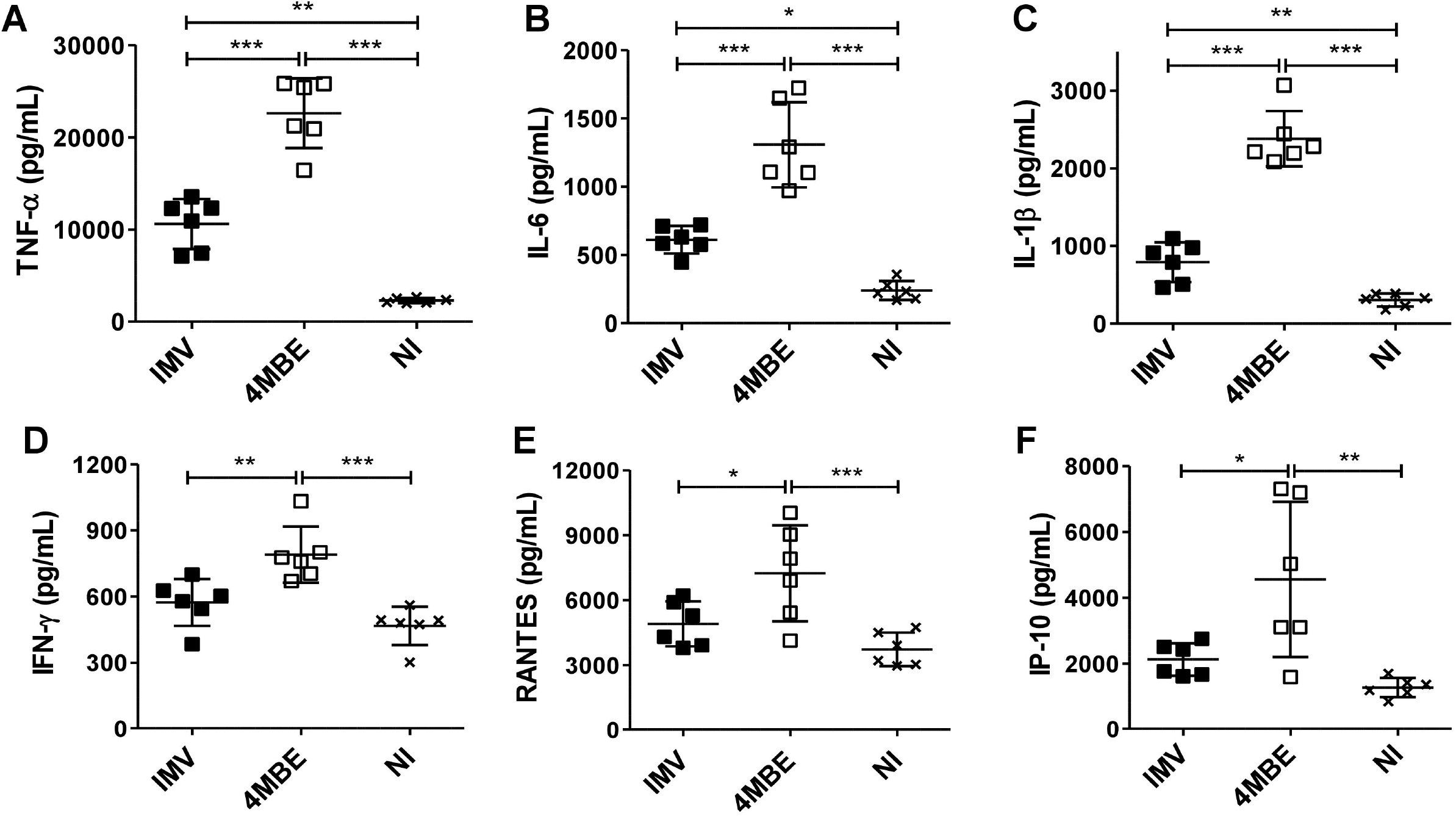
Macrophage inflammatory response is exacerbated upon infection by the 4MBE variant. Macrophages were infected by the IMV (black) or the 4MBE variant (white) at a MOI of 10:1 or not infected (NI, cross). (A) TNF-α, (B) IL-6, (C) IL-1β, (D) IFN-γ, (E) RANTES (CCL5) and (F) Interferon gamma-induced protein 10 (IP-10) release in cell culture supernatant was evaluated by Luminex^®^ Multiplex assay at 24 hpi. Values for each condition are the mean ± SD of three independent experiments, analyzed in duplicate. Means were compared using Repeated Measures ANOVA followed by Bonferroni correction.

It should be noted that both increased cell death and higher inflammatory response revealed by these bulk infection data could be, at least partially, related to the higher bacterial load upon infection by the 4MBE variant.

## DISCUSSION

The objective of the present study was to address the issue of within-host Mtb micro-diversity and its possible link with anti-TB treatment response. We therefore followed within-host evolution of Mtb clinical isolates in a cohort of patients with delayed culture conversion. We focused on minor variants detected by unfixed SNP, with a particular interest in drug tolerant sub-populations revealed after RIF exposure. The results support the absence of 2-month culture conversion as a predictive factor of poor outcome [20] as, among the 12 patients selected, 3 died and 8 required extended treatment duration in order to prevent failure. Interestingly, we were able to detect unfixed SNPs within the last Mtb isolates for 9/12 patients. When focusing on nsSNP, micro-diversity of Mtb clinical isolates was detected in 6/12 patients (equally distributed between early and last isolates). Although 8 additional unfixed nsSNP were revealed by RIF *in vitro* exposure, none was linked to drug resistance, consistent with previous reports on within-host Mtb evolution [10,12,16,31]. Of note, 3 unfixed SNP became fixed over the course of TB disease, while 2 other unfixed SNP became undetectable in the last Mtb isolate, consistent with previous observations that numerous variants emerge and then disappear during the course of infection and treatment within individual patients [16,31,32]. Interestingly, WGS of isolated colonies from a last Mtb isolate allowed detection of additional unfixed SNPs, previously undetected by WGS performed on the whole bacterial population, suggesting that more knowledge concerning Mtb diversity will gained in the future through the use of deeper sequencing.

Furthermore, we highlighted the plasticity of Mtb genetic micro-diversity of a clinical isolate, which presented a changing pattern of variant assembly depending on the experimental setting (intra-macrophage infection, antibiotic stress). Interestingly, a variant (4MBE variant) which was present at low frequency in the initial Mtb clinical isolate, became predominant under both stress conditions tested. Using a set of complementary approaches, we were able to show that this type of variant displayed tolerance to anti-TB treatment and improved intra-macrophagic persistence. We found that the 4MBE variant features a particular mycocerosates profile. Mycocerosates are components of PDIM and PGL, both playing important roles in Mtb host-pathogen interactions at many pathophysiological steps [23–25,33,34]. We also showed that the 4MBE variant has increased ability to infect macrophages, and to induce both macrophage inflammatory response and cell death (mainly apoptosis), possibly deleterious *in vivo* [28,30]. This pro-inflammatory profile could mainly favor infection initiation, and be less prone to develop in long-term infection, explaining the low frequency of the variant in the clinical isolate.

Current diagnostic tools are based on the detection of fixed or unfixed variants in loci involved in drug resistance, to adapt treatment as early as possible [35,36]. Moreover, features associated with Mtb tolerance have been suggested to be potential pre-resistant markers [37,38], as drug-tolerance and persistent infection may foster drug-resistance emergence [3]. Notwithstanding that C 3280555 G would be a hitchhiking SNP, this study shows that a minor variant may be associated with drug-tolerance and promote treatment failure as predicted by pharmacological modelling. A thorough study of such variants not necessarily linked to canonical drug-resistance, and their ability to promote anti-TB drugs tolerance may be crucial to prevent the subsequent emergence of resistance. Detecting unfixed antibiotic tolerant variants in Mtb clinical isolates, could thus provide new biomarkers to guide TB management in patients with treatment failure risk.

Interestingly, *in vitro* RIF exposure revealed the presence of a low-frequency variant within the Mtb bacterial population, otherwise undetected by the WGS bulk analysis. From an evolutionary perspective, given the lack of on-going horizontal gene exchange in Mtb, genetic micro-diversity, including minor variants with particular phenotypic profiles, could be envisioned as a pathogen feature required to effectively respond to changing environments, to persist, and to spread [16,31,32,39,40]. This emphasizes the importance of Mtb micro-diversity, which could become a useful biomarker to predict, in association with previously proven factors, which patients are at risk of poor response to anti-TB treatment. These patients may require personalized dosing and therapeutic management in order to improve the outcome and to effectively prevent treatment failure and resistance emergence.

## MATERIAL AND METHODS

### Study design

Patients were retrospectively enrolled among the 332 patients managed for drug-susceptible TB at Lyon university hospital between 2014 and 2016. Twelve patients with the following inclusion criteria were selected: positive *Mtb* culture at least 2 months after the beginning of anti-TB treatment and complete follow-up 2 years after treatment completion. Mtb isolates collection was approved by the *Comité de Protection des Personnes-Sud Est IV* (approval number: DC-2011-1306). All sampling and testing were performed for patient benefit. Written information was provided to patients and their families, complying with the French bioethics law.

### Bacterial strains

Mtb strains used in this study were isolated from pulmonary specimens during routine care in the Lyon University Hospital, France. The IMV and the 4MBE variant were obtained by cloning of the P1/L2 clinical isolate (Table 1). Clones were submitted to WGS to confirm the absence (IMV) or presence (4MBE) of the polymorphism of interest (C 3280555 G) and to confirm the absence of other mutations within the limits of the interpretation of short-read sequencing. Strains are stored at the Lyon University Hospital BSL3 laboratory and are available for research use if requested.

### TB-associated severity indices

#### The Bandim TB score

The modified Bandim TB score considers 5 symptoms (cough, hemoptysis, dyspnea, chest pain, night sweats) and 5 clinical findings (anemia, tachycardia, positive finding at lung auscultation, fever, BMI [<18 and <16]), with one point for each. One clinical finding was excluded, the mid upper arm circumference as this data was not available in the Lyon University Hospital. Accordingly, patients were stratified into two severity classes, mild (Bandim score ≤4) and moderate or severe (≥5) [41–43].

#### The MUST

The nutritional status of TB patients was evaluated thanks to the Malnutrition Universal Screening Tool (MUST), which includes three variables: unintentional weight loss score (weight loss < 5% = 0, weight loss 5-10% = 1, weight loss > 10% = 2), BMI score (BMI > 20.0 = 0, BMI 18.5-20.0 = 1, BMI < 18.5 = 2) and anorexia (if yes = 2). A MUST ≥ 4 is associated with poor TB prognosis [44].

#### MIC determination

Minimum inhibitory concentrations (MIC) were determined for RIF and INH (Sigma-Aldrich, Saint Louis, MO, USA) by a standard microdilution method as previously described [45]. Briefly, the assay was performed in a 96-well microtiter plate, with the concentrations ranging from 8 to 0.0008 mg/L, with a final inoculum of approximately 1.10^5^ CFU/mL in each well. The plates were incubated at 37° C for 12 to 18 days, and the wells were assessed for visible turbidity. The lowest concentration at which there was no visible turbidity was defined as the MIC.

#### Antibiotics exposure using the BACTEC 960 system

Antibiotic exposure experiments were performed in Mycobacterial Growth Indicator Tube (MGIT) using the BACTEC 960 system (Becton Dickinson, Sparks, MD, USA). Antibiotic solutions were added to MGIT at the required concentration and inoculated with 10^5^ CFU/mL. For the drug-free growth control, MGIT were inoculated with 10^3^ CFU/mL. Analysis of the fluorescence was used to determine whether the tube was instrument positive; i.e. the test sample contains viable organisms, and results were expressed using MGIT time to positivity (TTP) system, reflecting bacterial growth [46]. Bacterial cultures were recovered to proceed to DNA or RNA extraction and purification.

#### Mtb RNA extraction and reverse transcription

Mycobacteria were incubated with RNAprotect Bacteria Reagent (Qiagen, Valencia, CA, USA), lysed and Mtb total RNA was isolated by using RNeasy Plus mini kit according to the manufacturer’s instruction (Qiagen). Reverse transcription was performed by using the Reverse Transcription System according to the manufacturer’s instruction (Promega, Fitchburg, WI, USA).

#### Droplet digital PCR

ddPCR procedure was conducted using QX100 Droplet Digital PCR System (Bio-Rad Laboratories, Hercules, CA, USA). Total reaction volume was 22 μL, containing 11 μL of 2X ddPCR Supermix for Probes (No dUTPs), 8.9 μL of template DNA, 10 U of *Hind*III restriction enzyme and 1.1 μL of a 20X primers-probes mix (ddPCR mutant assay) provided by Bio-Rad (concentrations and sequences of probes are not provided by the manufacturer but they guaranty the validation by Minimum Information for Publication of Quantitative Real-Time PCR Experiments [MIQE] guidelines). The sequence studied was CGACACCGTTCGTGACCTCATCGCCCGTTGGGAGCAGCGGGACGTGATGGCGCGC GAGGTG[G/C]CCGTCGACGTGGCGTCGCACTCGCCTCAAGTCGATCCGATACTCGACGATTTGGCCGCGGC and the product was 70 bp. A no-template control and a positive control were used in each ddPCR batch. The procedure was carried out with droplet generation then PCR amplification with the following conditions: 95°C for 10 min followed by 40 cycles of 94°C for 30 s and 55°C for 1 min, and 98°C for 10 min, all steps with a ramp rate of 2°C/s. Analysis were done with QuantaSoft Analysis Pro software version 1.0.596 (Bio-Rad Laboratories). Positive droplets were counted and percentages of wild-type and mutant templates were calculated (% mutant =[(n mixed population/2)+n mutant]/total n amplified).

#### DNA extraction, targeted NGS and WGS

Genomic DNA was purified from cleared lysate using a QIAamp DNA mini Kit (Qiagen). For targeted NGS, we performed PCR using Platinum SuperFi DNA polymerase (Life Technologies SAS, Carlsbad, CA, USA) and primers indicated in S3 Table. DNA libraries were prepared with Nextera XT kit (Illumina, San Diego, CA, USA). Samples were sequenced on NextSeq or MiSeq system (Illumina) to produce 150 or 300 base-pair paired-end reads at the Bio-Genet NGS facility of Lyon University Hospital, as previously described [47].

#### Bioinformatic analysis of Illumina data

Reads were mapped with BOWTIE2 to the Mtb H37Rv reference genome (Genbank NC000962.2) and variant calling was made with SAMtools mpileup, as previously described [47]. A valid nucleotide variant was called if the position was covered by a depth of at least 10 reads and supported by a minimum threshold rate of 10%. Regions with repetitive or similar sequences were excluded, i.e. PE, PPE, PKS, PPS, ESX. The WGS reference genome coverage ranges 96.5% to 99.0%, with an average depth of coverage of 46x to 185x.

Sequences have been submitted to European Nucleotide Archive (ENA) under accession number PRJEB37306.

#### Apolar lipid extraction and analysis by gas chromatography–mass spectrometry (GC-MS)

Mycobacteria were grown in Middlebrooks 7H9 medium supplemented with 10% OADC (oleic acid, albumin, dextrose, catalase) until mid-log phase and pelleted before being washed twice in sterile water. The pellet was resuspended in methanol: 0.3% aqueous NaCl (10:1) and petroleum ether was added to separate the liquid layers. The non-aqueous petroleum ether extracts containing apolar lipids were dried under nitrogen. Apolar lipids were hydrolyzed with KOH 5% in methanol at 100°C for 90 min. The reaction mixture was acidified with 3N HCl and acidic water was added. Fatty acids released were extracted with chloroform-ethanol (3:1). The upper phase was removed and lower phase stored. The extracted fatty acids were derivatized with pentafluorobenzyl bromide (PFB) and analyzed in the negative ion mode by gas chromatography-mass spectrometry (GC-MS/MS Agilent HP7890B/7000C). The gas chromatograph was performed at the Functional Lipidomics platform, acknowledged by Infrastructure in Biology, Health and Agronomy (IBiSA).

#### Killing kinetics of RIF and INH alone and combination and mathematical modeling

*In vitro* experiments of the antibacterial effect of INH and RIF alone and in combination, as well as the mathematical modeling of the results were performed as described in a previous publication [45].

##### Killing kinetics of single drug

For each strain, an inoculum of approximately 5.10^5^ CFU/mL was prepared in 7H9 Middlebrook medium. The inoculum was incubated without any antibiotic (growth control) and with RIF, at concentrations of 1/32, 1/16, 1/8, 1/4, 1/2, 1, 2, 4, 16, 64, and 256 multiples of MIC (xMIC). Incubation was performed in 96 deepwell plates (1 ml working volume) at 37°C. Each well was homogenized by pipetting and vortexing, serially diluted, and plated on 7H10 agar. The numbers of CFU per milliliter was determined on alternate days after 3, 5, 7, and 10 days of drug exposure. Each experiment was performed twice.

##### Killing kinetics of INH + RIF combination

For each strain, a bacterial inoculum was prepared as described above. Twelve concentrations of IHN (0, 1/32, 1/16, 1/8, 1/4, 1/2, 1, 2, 4, 8, 16, and 64 multiples of MIC) were combined with 12 possible concentrations of RIF (same multiples of MIC as INH) in an incomplete checkerboard design. The incubation was performed as for single drug experiments. The numbers of CFU per milliliter was determined after day 7 of drug exposure. A total of 142 bacterial counts were available for each strain.

##### Mathematical modelling of single drug effect

A Hill pharmacodynamic equation was used to describe the antibacterial effect of single drugs [48]. The model was fit to single drug log-transformed data, as follows:

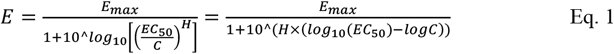

Where E is the drug effect, C is the drug concentration, E_max_ is the maximal effect EC_50_ is the median effect concentration, and H is the Hill coefficient of sigmoidicity. GraphPad Prism^®^ for Windows version 5.02 (GraphPad Software, La Jolla, CA, USA) was used for model fitting and parameter estimation.

The antibacterial effect E considered was the net difference between the bacterial count measured with no drug (C = 0) and the count for a given drug concentration at the same time-point. Drug concentrations were normalized to the MIC, so C is expressed as the ratio of the actual concentration of drug in culture divided by the MIC.

##### Response-surface modeling of INH and RIF combined effect

We used the Minto model [45,49] to describe the combined antimicrobial effect of INH and RIF based on data measured after 7 days of therapy.

The Minto model is an extension of the Hill equation to drug combination and is defined as follows as follows:

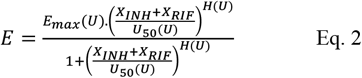

With 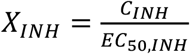 and 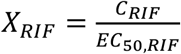

And 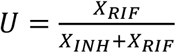

In those equations, X_INH_ and X_RIF_ are concentrations of INH and RIF normalized to the drug potency, U quantifies the ratio of each drug in the combination, E_max_(U) is the maximal effect at ratio U, U_50_(U) is the number of units of X_RIF_ associated with 50% of the maximal effect at ratio U, and H(U) is the coefficient of sigmoidicity at ratio U. Each of the three parameters of the Hill equation are function of U, the drug ratio. The concentration-effect curve associated with each ratio defined the contour of a response-surface of the drug combination.

As suggested by Minto et al., we used a polynomial function to describe E_max_(U), U_50_(U), and H(U), as described below:

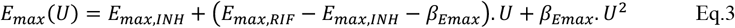

where E_max,INH_ and E_max,RIF_ are the maximal effect of INH and RIF alone, respectively, and β_Emax_ is the coefficient of the two-order polynomial for E_max_(U);

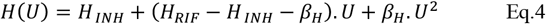

where H_INH_ and H_RIF_ are the Hill coefficient of sigmoidicity for INH and RIF alone, respectively, and β_H_ is the coefficient of the two-order polynomial for H(U); and

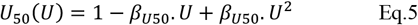

Where β_U50_ is the coefficient of the two-order polynomial for U_50_(U).

The last equation allows to interpret the response surface in terms of synergy/antagonism. If β_U50_ is not different from 0, U_50_ is equal to 1 for all values of U, and the interaction is additive. If β_U50_ is significantly greater than 0, the curve shows an inward curvature and the normalized drug mixture, is more than additive, which means synergy. If β_U50_ is significantly lower than 0, the curve displays an outward curvature, and the normalized drug mixture (X_INH_ + X_RIF_)/U_50_(U)) is less than additive, which means antagonism.

The Minto model was fit to data by using non-linear regression within the Matlab software (version 2011b; The Mathworks, Natick, MA, USA). Point estimates of model parameters along with their confidence intervals were obtained. Goodness-of-fit of the model was assessed by analysis of plots of observed versus predicted antibacterial effects

Values and 95% confidence intervals of the β_U50_ coefficient were examined to interpret the combined action in terms of synergy/antagonism. Synergy was confirmed when the lower bound of the 95% confidence interval was greater than 0. Antagonism was stated when the upper bound of the 95% confidence interval was lower than 0. When the 95% confidence interval included 0, the interaction was considered as additive.

#### Cell Culture and Infection protocol

U937 cells, monocytic cell line from pleural effusion, cultured in RPMI-1640 medium supplemented with 10% fetal bovine serum were differentiated with phorbol 12-myristate 13-acetate (PMA; Sigma-Aldrich) 100nM for 48h at a density of 2.10^5^ cells/mL. Adherent cells were then infected with bacterial suspension at a multiplicity of infection (MOI) of 10:1 (bacteria to cells) in antibiotic-free culture medium, at 37 °C in 5% CO_2_ atmosphere, as previously described [50].

#### Intracellular CFU counting

U937 cells differentiated cells were infected by Mtb, as described above. After 5 h of culture, infected cells were incubated with amikacin 200 mg/L for 1 h to remove the extracellular Mtb. The cells were washed thrice to remove amikacin. To study invasiveness, cells were immediately lysed with distilled water containing saponin 0.1% for 10 min and intracellular mycobacteria were plated on complete 7H10 agar and incubated for 3-4 weeks. To measure intra-cellular multiplication, separate wells with amikacin-treated inoculated cells were further incubated for 96 h at 37 °C in 5 % CO_2_ atmosphere. After that, extracellular bacteria were removed by 1 h of incubation with amikacin 200 mg/L and intracellular bacteria were counted as describe above.

#### Immunofluorescence and image acquisition

U937 differentiated cells were infected by Mtb, as described above, before staining for confocal microscopy analysis, 6 h and 24 h post-infection (hpi), as previously described [50]. For the study of autophagic flux, 3 h before staining, cells were incubated with chloroquine (Sigma-Aldrich) 40μM to avoid acidification of autophagy compartments. Infected cells were washed thrice and incubated with LysoTracker Red DND-99 500nM (Thermo Fisher Scientific Inc, Rockford, IL, US; L7528) for 1 h to detect acidify compartments. Following 4% paraformaldehyde fixation over 20 min, cells were incubated with 100 mM glycine for 20 min and washed with PBS. Cells were then permeabilized with 0.1% Triton X-100 for 10 min and saturated in PBS-3% bovine serum albumin (BSA) for 30 min. Cells were labelled with anti-Mtb coupled FITC antibody (Abcam, Cambridge, MA, US; ab20962, 1/100 dilution) and anti-LC3B coupled Dy-Light 650 antibody (to assess autophagy activation; Thermo Fisher Scientific, PA5-22937, 1/400 dilution) in PBS-3% BSA-0.05% Triton X100 for 1h at room temperature. Slides were mounted in Fluoromount medium (Sigma-Aldrich) and observed with a Leica confocal microscope SP5X. Images were acquired with Leica Application Suite software. Stacks of confocal images and quantification of percent of colocalization were performed with ImageJ software [51] and JACoP plugin.

#### CCF-4 assay for translocation of Mtb in host cell cytosol

To detect mycobacterial escape from the phagosome and translocation in host cell cytosol, the CCF4 FRET assay was performed [24,52]. Briefly, infected cells were stained with 50 μM CCF4 (Invitrogen, Carlsbad, CA, USA) in buffer containing anion transport inhibitor (Invitrogen) for 2 h at room temperature, as recommended by the manufacturer. Cells were washed with PBS containing anion transport inhibitor before fixing with paraformaldehyde (PFA, Sigma-Aldrich) 4% for 30 min at room temperature in the dark. Cells were washed before performing directly fluorescence acquisition. Intact CCF4-AM emits green fluorescence (520 nm) due to FRET between the fluorescent moieties, indicating that mycobacteria are in intracellular compartments. Cleavage of CCF4-AM, due to β-lactamase expressed by cytosolic Mtb, leads to a shift in the fluorescence emitted to bleu (450nm). The results obtained were normalized on intracellular bacterial load.

#### Measurement of lactate dehydrogenase (LDH) in cell culture supernatants

U937 cells differentiated cells were infected by Mtb, as described above. To quantify necrosis of U937 cells, the LDH release in cell culture supernatant from infected cells was measured at 24 h and 96 hpi by using the colorimetric kit (Roche, Mannheim, Germany) and following the manufacturer’s instructions.

#### Quantitative determination of apoptosis

U937 differentiated cells were infected by Mtb, as described above, before staining for cell death analysis, at 24 hpi and 96 hpi. Infected cells were washed thrice and apoptosis was detected by Annexin V FITC (Thermo Fisher Scientific) after 15min of incubation at room temperature. After fixing with PFA 4% for 30min, cells were analyzed by using fluorescent microscopy.

#### Infected macrophage RNA extraction

U937 differentiated cells were infected by Mtb, as described above, before RNA extraction. Total macrophage RNA was obtained from Mtb-infected cells at 24 hpi using the Trizol method as previously described [53]. Briefly, macrophages were lysed with Trizol reagent (Qiagen) and total RNA was isolated with miRNeasy mini kit according to the manufacturer’s instruction (Qiagen). The macrophage RNA was subjected to DNaseI digestion before final purification.

#### Transcriptomic analysis

Transcriptomic analysis was performed at ViroScan3D/ProfileXpert platform (www.viroscan3d.com). Briefly, RNA from each sample was quantified using Quantus HSRNA method (Promega) and qualified using Fragment analyzer HSRNA (AATI). After quality control, total RNA were submitted to polyA capture using NextFlex poly(A) Beads (PerkinElmer, Waltham, MA, USA), then mRNA were submitted to library preparation using NextFlex Rapid Directional mRNA (PerkinElmer) and 100 ng input. As recommended by the ENCODE (Encyclopedia of DNA Elements) consortium ERCC (External RNA Control Consortium) RNA spike-In (Invitrogen) were added to samples in order to ensure reproducibility of the experiments. Single-read sequencing with 75 bp read length was performed on NextSeq 500 high output flowcell (Illumina). Mapped reads for each samples were counted and normalized using FPKM method [54]. Fold change between the different groups were calculated using median of groups and p-value of difference were calculated using t-test with equal variance without p-value correction in the RStudio v.0.99.893 (RStudio Inc., Boston, MA, USA). Heatmaps were performed thanks to Heatmapper [55], using average linkage clustering method and Pearson distance measurement method.

#### Cytokine production assay

U937 differentiated cells were infected by Mtb, as described above, and cell supernatant was recovered 6 hpi, 24 hpi and 96 hpi. Cell culture supernatants were screened for the presence of 27 human cytokines and chemokines using the Bio-Plex Pro Human Cytokine Standard 27-Plex kit (Bio-Rad Laboratory) on a FLEXMAP 3D^®^ analyzer (Luminex, Austin, TX, USA). Data were analyzed using Bio-Plex Manager software version 6.1 (Bio-Rad Laboratory).

#### Statistical analysis

Statistical analyses were performed using GraphPad Prism for Windows, version 5.02 (GraphPad Software, La Jolla, CA, USA). Data obtained from lipidomic analysis, RIF exposure experiments, variant frequency analysis by ddPCR, intracellular CFU counting, lytic cell death, as well as cytokine and chemokine production were expressed as mean ± standard deviation (SD). Means were compared using One-Way ANOVA or Repeated Measures ANOVA followed with Bonferroni correction, unpaired t-test or Student’s t-test, where appropriate. For data obtained from pharmacological models, parameter values were given as point estimate [95% confidence interval, CI)]. For confocal microscopy analysis and macrophage apoptosis, data were expressed as median values [interquartile range, IQR]. Results were compared by using Kruskal-Wallis analysis, using Dunn’s Multiple Comparison Test or Mann Whitney test, where appropriate. \**p*<0.05, ** *p* <0.01, \*\*\**p* <0.001.

## ACKNOWLEDGMENTS

The authors thank Philip Robinson and Véréna Landel (DRCI, Hospices Civils de Lyon) for help in manuscript preparation.

## Author Contributions

Conceptualization: C.Ge., S.G., O.D.; Methodology: C.Ge., C.Gi., S.G., O.D.; Software: C.Ge., J-L.B., E.W.; Investigation and validation: C.Ge., E.H., A.B., J.H., F.B., M.G.; Data analysis: C.Ge., A.B., E.H., F.B., M.G., C.Gi., S.G., O.D.; Data interpretation: C.Ge., A.B., E.H., J-L.B., J.H., S.V., C.Gi., S.G., O.D.; Visualization: C.Ge, E.H., S.G., O.D.; Writing – Original Draft: C.Ge., S.G., O.D.; Writing – Review & Editing: All authors; Funding Acquisition: C.Ge., J-L.B., S.V., G.L., S.G., O.D.

## SUPPORTING INFORMATION

**S1 Fig. Improved Mtb intra-macrophagic survival of the 4MBE variant.** At 24 hpi, cells were stained with anti-Mtb coupled FITC. (A) The proportion of infected cells were determined across at least 10 confocal images, containing between 30 and 80 cells per image, per replicate. (B) Mtb staining area was determined across at least 40 cells per replicate. Values for each condition are the median values [interquartile range, IQR] of at least three independent experiments. Statistical significance was determined using Mann Whitney test.

**S2 Fig. Early inflammatory response is slightly exacerbated upon infection by the 4MBE variant.** (A) TNF-α, (B) IL-6, (C) IL-1β, (D) IFN-γ, (E) CCL5 or RANTES and (F) Interferon gamma-induced protein 10 (IP-10) release in cell culture supernatant was evaluated at 6 hpi. Values for each condition are the mean ± standard deviation (SD) of three independent experiments. Means were compared using Repeated Measures ANOVA followed by Bonferroni correction.

**S3 Fig. Persistent inflammatory response upon infection by the 4MBE variant.** (A) TNF-α, (B) IL-6, (C) IL-1β, (D) IFN-γ, (E) CCL5 or RANTES and (F) Interferon gamma-induced protein 10 (IP-10) release in cell culture supernatant was evaluated at 96 hpi. Values for each condition are the mean ± standard deviation (SD) of three independent experiments. Means were compared using Repeated Measures ANOVA followed by Bonferroni correction.

**S1 Table. Model parameters and fit for rifampicin (RIF) effect measured after 7 days.**

**S2 Table. Parameter values and goodness of fit of the response-surface model describing the combined action of INH and RIF measured after 7 days.**

**S3 Table. Primer sets for targeted NGS.**

## REFERENCES

1. World Health Organization, Geneva | Global tuberculosis report 2019. WHO 2019. Available: http://www.who.int/tb/publications/global_report/en/

2. Wallis RS, Patil S, Cheon S-H, Edmonds K, Phillips M, Perkins MD, et al. Drug Tolerance in *Mycobacterium tuberculosis*. Antimicrob Agents Chemother. 1999;43: 2600–2606.

3. Levin-Reisman I, Ronin I, Gefen O, Braniss I, Shoresh N, Balaban NQ. Antibiotic tolerance facilitates the evolution of resistance. Science. 2017;355: 826–830. doi:10.1126/science.aaj2191

4. Brown AC, Bryant JM, Einer-Jensen K, Holdstock J, Houniet DT, Chan JZM, et al. Rapid Whole-Genome Sequencing of *Mycobacterium tuberculosis* Isolates Directly from Clinical Samples. Land GA, editor. J Clin Microbiol. 2015;53: 2230–2237. doi:10.1128/JCM.00486-15

5. Casali N, Broda A, Harris SR, Parkhill J, Brown T, Drobniewski F. Whole Genome Sequence Analysis of a Large Isoniazid-Resistant Tuberculosis Outbreak in London: A Retrospective Observational Study. PLoS Med. 2016;13. doi:10.1371/journal.pmed.1002137

6. Doyle RM, Burgess C, Williams R, Gorton R, Booth H, Brown J, et al. Direct Whole-Genome Sequencing of Sputum Accurately Identifies Drug-Resistant *Mycobacterium tuberculosis* Faster than MGIT Culture Sequencing. Mellmann A, editor. J Clin Microbiol. 2018;56. doi:10.1128/JCM.00666-18

7. Genestet C, Hodille E, Westeel E, Ginevra C, Ader F, Venner S, et al. Subcultured *Mycobacterium tuberculosis* isolates on different growth media are fully representative of bacteria within clinical samples. Tuberc Edinb Scotl. 2019;116: 61–66. doi:10.1016/j.tube.2019.05.001

8. Shockey AC, Dabney J, Pepperell CS. Effects of Host, Sample, and *in vitro* Culture on Genomic Diversity of Pathogenic Mycobacteria. Front Genet. 2019;10. doi:10.3389/fgene.2019.00477

9. Votintseva AA, Bradley P, Pankhurst L, del Ojo Elias C, Loose M, Nilgiriwala K, et al. Same-Day Diagnostic and Surveillance Data for Tuberculosis via Whole-Genome Sequencing of Direct Respiratory Samples. Tang Y-W, editor. J Clin Microbiol. 2017;55: 1285–1298. doi:10.1128/JCM.02483-16

10. Nimmo C, Brien K, Millard J, Grant AD, Padayatchi N, Pym AS, et al. Dynamics of within-host *Mycobacterium tuberculosis* diversity and heteroresistance during treatment. EBioMedicine. 2020;55: 102747. doi:10.1016/j.ebiom.2020.102747

11. Lieberman TD, Wilson D, Misra R, Xiong LL, Moodley P, Cohen T, et al. Genomic diversity in autopsy samples reveals within-host dissemination of HIV-associated *Mycobacterium tuberculosis*. Nat Med. 2016;22: 1470–1474. doi:10.1038/nm.4205

12. O’Neill MB, Mortimer TD, Pepperell CS. Diversity of *Mycobacterium tuberculosis* across Evolutionary Scales. PLOS Pathog. 2015;11: e1005257. doi:10.1371/journal.ppat.1005257

13. Pérez-Lago L, Comas I, Navarro Y, González-Candelas F, Herranz M, Bouza E, et al. Whole genome sequencing analysis of intrapatient microevolution in *Mycobacterium tuberculosis*: potential impact on the inference of tuberculosis transmission. J Infect Dis. 2014;209: 98–108. doi:10.1093/infdis/jit439

14. Vargas R, Freschi L, Marin M, Epperson LE, Smith M, Oussenko I, et al. In-host population dynamics of *M. tuberculosis* during treatment failure. bioRxiv. 2019; 726430. doi:10.1101/726430

15. Copin R, Wang X, Louie E, Escuyer V, Coscolla M, Gagneux S, et al. Within Host Evolution Selects for a Dominant Genotype of *Mycobacterium tuberculosis* while T Cells Increase Pathogen Genetic Diversity. PLOS Pathog. 2016;12: e1006111. doi:10.1371/journal.ppat.1006111

16. Ley SD, de Vos M, Van Rie A, Warren RM. Deciphering Within-Host Microevolution of *Mycobacterium tuberculosis* through Whole-Genome Sequencing: the Phenotypic Impact and Way Forward. Microbiol Mol Biol Rev MMBR. 2019;83. doi:10.1128/MMBR.00062-18

17. Navarro Y, Pérez-Lago L, Sislema F, Herranz M, de Juan L, Bouza E, et al. Unmasking subtle differences in the infectivity of microevolved *Mycobacterium tuberculosis* variants coinfecting the same patient. Int J Med Microbiol IJMM. 2013;303: 693–696. doi:10.1016/j.ijmm.2013.10.002

18. Azarian T, Ridgway JP, Yin Z, David MZ. Long-Term Intrahost Evolution of Methicillin Resistant *Staphylococcus aureus* Among Cystic Fibrosis Patients With Respiratory Carriage. Front Genet. 2019;10. doi:10.3389/fgene.2019.00546

19. Winstanley C, O’Brien S, Brockhurst MA. *Pseudomonas aeruginosa* Evolutionary Adaptation and Diversification in Cystic Fibrosis Chronic Lung Infections. Trends Microbiol. 2016;24: 327–337. doi:10.1016/j.tim.2016.01.008

20. Mitchison DA. Assessment of New Sterilizing Drugs for Treating Pulmonary Tuberculosis by Culture at 2 Months. Am Rev Respir Dis. 1993;147: 1062–1063. doi:10.1164/ajrccm/147.4.1062

21. Herbst DA, Jakob RP, Zähringer F, Maier T. Mycocerosic acid synthase exemplifies the architecture of reducing polyketide synthases. Nature. 2016;531: 533–537. doi:10.1038/nature16993

22. Trivedi OA, Arora P, Vats A, Ansari MZ, Tickoo R, Sridharan V, et al. Dissecting the mechanism and assembly of a complex virulence mycobacterial lipid. Mol Cell. 2005;17: 631–643. doi:10.1016/j.molcel.2005.02.009

23. Ferreras JA, Stirrett KL, Lu X, Ryu J-S, Soll CE, Tan DS, et al. Mycobacterial Phenolic Glycolipid Virulence Factor Biosynthesis: Mechanism and Small-Molecule Inhibition of Polyketide Chain Initiation. Chem Biol. 2008;15: 51–61. doi:10.1016/j.chembiol.2007.11.010

24. Quigley J, Hughitt VK, Velikovsky CA, Mariuzza RA, El-Sayed NM, Briken V. The Cell Wall Lipid PDIM Contributes to Phagosomal Escape and Host Cell Exit of *Mycobacterium tuberculosis*. mBio. 2017;8: e00148–17. doi:10.1128/mBio.00148-17

25. Siegrist MS, Bertozzi CR. Mycobacterial Lipid Logic. Cell Host Microbe. 2014;15: 1–2. doi:10.1016/j.chom.2013.12.005

26. Nicoara SC, Minnikin DE, Lee OCY, O’Sullivan DM, McNerney R, Pillinger CT, et al. Development and optimization of a gas chromatography/mass spectrometry method for the analysis of thermochemolytic degradation products of phthiocerol dimycocerosate waxes found in *Mycobacterium tuberculosis*. Rapid Commun Mass Spectrom RCM. 2013;27: 2374–2382. doi:10.1002/rcm.6694

27. Redman JE, Shaw MJ, Mallet AI, Santos AL, Roberts CA, Gernaey AM, et al. Mycocerosic acid biomarkers for the diagnosis of tuberculosis in the Coimbra Skeletal Collection. Tuberc Edinb Scotl. 2009;89: 267–277. doi:10.1016/j.tube.2009.04.001

28. Parandhaman DK, Narayanan S. Cell death paradigms in the pathogenesis of *Mycobacterium tuberculosis* infection. Front Cell Infect Microbiol. 2014;4. doi:10.3389/fcimb.2014.00031

29. Teng O, Ang CKE, Guan XL. Macrophage–Bacteria Interactions—A Lipid-Centric Relationship. Front Immunol. 2017;8. doi:10.3389/fimmu.2017.01836

30. Zhai W, Wu F, Zhang Y, Fu Y, Liu Z. The Immune Escape Mechanisms of *Mycobacterium Tuberculosis*. Int J Mol Sci. 2019;20. doi:10.3390/ijms20020340

31. Black PA, de Vos M, Louw GE, van der Merwe RG, Dippenaar A, Streicher EM, et al. Whole genome sequencing reveals genomic heterogeneity and antibiotic purification in *Mycobacterium tuberculosis* isolates. BMC Genomics. 2015;16: 857. doi:10.1186/s12864-015-2067-2

32. Trauner A, Liu Q, Via LE, Liu X, Ruan X, Liang L, et al. The within-host population dynamics of *Mycobacterium tuberculosis* vary with treatment efficacy. Genome Biol. 2017;18: 71. doi:10.1186/s13059-017-1196-0

33. Jackson M. The Mycobacterial Cell Envelope—Lipids. Cold Spring Harb Perspect Med. 2014;4. doi:10.1101/cshperspect.a021105

34. Queval CJ, Brosch R, Simeone R. The Macrophage: A Disputed Fortress in the Battle against *Mycobacterium tuberculosis*. Front Microbiol. 2017;8. doi:10.3389/fmicb.2017.02284

35. Ng KCS, Supply P, Cobelens FGJ, Gaudin C, Gonzalez-Martin J, Jong BC de, et al. How Well Do Routine Molecular Diagnostics Detect Rifampin Heteroresistance in *Mycobacterium tuberculosis?* J Clin Microbiol. 2019;57: e00717–19. doi:10.1128/JCM.00717-19

36. Sun G, Luo T, Yang C, Dong X, Li J, Zhu Y, et al. Dynamic population changes in *Mycobacterium tuberculosis* during acquisition and fixation of drug resistance in patients. J Infect Dis. 2012;206: 1724–1733. doi:10.1093/infdis/jis601

37. Pasipanodya JG, Gumbo T. A new evolutionary and pharmacokinetic–pharmacodynamic scenario for rapid emergence of resistance to single and multiple anti-tuberculosis drugs. Curr Opin Pharmacol. 2011;11: 457–463. doi:10.1016/j.coph.2011.07.001

38. Schmalstieg AM, Srivastava S, Belkaya S, Deshpande D, Meek C, Leff R, et al. The Antibiotic Resistance Arrow of Time: Efflux Pump Induction Is a General First Step in the Evolution of Mycobacterial Drug Resistance. Antimicrob Agents Chemother. 2012;56: 4806–4815. doi:10.1128/AAC.05546-11

39. Buckling A, Brockhurst MA. Kin selection and the evolution of virulence. Heredity. 2008;100: 484–488. doi:10.1038/sj.hdy.6801093

40. Nair RR, Sharan D, Ajitkumar P. A Minor Subpopulation of Mycobacteria Inherently Produces High Levels of Reactive Oxygen Species That Generate Antibiotic Resisters at High Frequency From Itself and Enhance Resister Generation From Its Major Kin Subpopulation. Front Microbiol. 2019;10. doi:10.3389/fmicb.2019.01842

41. Dewi DNSS, Mertaniasih NM, Soedarsono. Severity Of TB Classified By Modified Bandim TB Scoring Associates With The Specific Sequence Of EsxA Genes In MDR-TB Patients. Afr J Infect Dis. 2020;14: 8–15. doi:10.21010/ajid.v14i1.2

42. Rudolf F, Joaquim LC, Vieira C, Bjerregaard-Andersen M, Andersen A, Erlandsen M, et al. The Bandim tuberculosis score: reliability and comparison with the Karnofsky performance score. Scand J Infect Dis. 2013;45: 256–264. doi:10.3109/00365548.2012.731077

43. Wejse C, Gustafson P, Nielsen J, Gomes VF, Aaby P, Andersen PL, et al. TBscore: Signs and symptoms from tuberculosis patients in a low-resource setting have predictive value and may be used to assess clinical course. Scand J Infect Dis. 2008;40: 111–120. doi:10.1080/00365540701558698

44. Miyata S, Tanaka M, Ihaku D. The prognostic significance of nutritional status using malnutrition universal screening tool in patients with pulmonary tuberculosis. Nutr J. 2013;12: 42. doi:10.1186/1475-2891-12-42

45. Genestet C, Ader F, Pichat C, Lina G, Dumitrescu O, Goutelle S, et al. Combined Antibacterial Effect of Isoniazid and Rifampicin on Four *Mycobacterium tuberculosis* Strains: in Vitro Experiments and Response-Surface Modeling. Antimicrob Agents Chemother. 2017. doi:10.1128/AAC.01413-17

46. Bowness R, Boeree MJ, Aarnoutse R, Dawson R, Diacon A, Mangu C, et al. The relationship between *Mycobacterium tuberculosis* MGIT time to positivity and CFU in sputum samples demonstrates changing bacterial phenotypes potentially reflecting the impact of chemotherapy on critical sub-populations. J Antimicrob Chemother. 2015;70: 448–455. doi:10.1093/jac/dku415

47. Genestet C, Tatai C, Berland J-L, Claude J-B, Westeel E, Hodille E, et al. Prospective Whole-Genome Sequencing in Tuberculosis Outbreak Investigation, France, 2017–2018. Emerg Infect Dis. 2019;25: 589–592. doi:10.3201/eid2503.181124

48. Goutelle S, Maurin M, Rougier F, Barbaut X, Bourguignon L, Ducher M, et al. The Hill equation: a review of its capabilities in pharmacological modelling. Fundam Clin Pharmacol. 2008;22: 633–648. doi:10.1111/j.1472-8206.2008.00633.x

49. Minto CF, Schnider TW, Short TG, Gregg KM, Gentilini A, Shafer SL. Response Surface Model for Anesthetic Drug Interactions. J Am Soc Anesthesiol. 2000;92: 1603–1616.

50. Genestet C, Bernard-Barret F, Hodille E, Ginevra C, Ader F, Goutelle S, et al. Antituberculous drugs modulate bacterial phagolysosome avoidance and autophagy in *Mycobacterium tuberculosis-infected* macrophages. Tuberc Edinb Scotl. 2018;111: 67–70. doi:10.1016/j.tube.2018.05.014

51. Schneider CA, Rasband WS, Eliceiri KW. NIH Image to ImageJ: 25 years of image analysis. Nat Methods. 2012;9: 671–675. doi:10.1038/nmeth.2089

52. Simeone R, Bobard A, Lippmann J, Bitter W, Majlessi L, Brosch R, et al. Phagosomal Rupture by *Mycobacterium tuberculosis* Results in Toxicity and Host Cell Death. PLOS Pathog. 2012;8: e1002507. doi:10.1371/journal.ppat.1002507

53. Koo M-S, Subbian S, Kaplan G. Strain specific transcriptional response in *Mycobacterium tuberculosis* infected macrophages. Cell Commun Signal CCS. 2012;10: 2. doi:10.1186/1478-811X-10-2

54. Trapnell C, Williams BA, Pertea G, Mortazavi A, Kwan G, van Baren MJ, et al. Transcript assembly and abundance estimation from RNA-Seq reveals thousands of new transcripts and switching among isoforms. Nat Biotechnol. 2010;28: 511–515. doi:10.1038/nbt.1621

55. Babicki S, Arndt D, Marcu A, Liang Y, Grant JR, Maciejewski A, et al. Heatmapper: web-enabled heat mapping for all. Nucleic Acids Res. 2016;44: W147–W153. doi:10.1093/nar/gkw419

