## Supplementary material for "Rifampicin exposure reveals within-host *Mycobacterium tuberculosis* diversity in patients with delayed culture conversion": Table S1

**S1 Table. Model parameters and fit for rifampicin (RIF) effect measured after 7 days**

|  | ***Mycobacterium tuberculosis* strains** | |
| --- | --- | --- |
| **Parameter** | **IMV** | **4MBE** |
| E_max_ (decline in log_10_ CFU/mL) | 7.1 [6.6; 7.7] | 6.9 [6.1; 7.7] |
| EC_50_ (x MIC) | 1.0 [0.6; 1.6] | 5.9 [3.4; 10.4] |
| Hill coefficient (H) | 0.49 [0.41; 0.58] | 0.61 [0.49; 0.74] |
| Model fit (R^2^) | 0.98 | 0.98 |

Parameter values are given as point estimate with [95% confidence interval], except for R².

MIC: minimum inhibitory concentration.
