## Supplementary material for "Rifampicin exposure reveals within-host *Mycobacterium tuberculosis* diversity in patients with delayed culture conversion": Table S2

**S2 Table. Parameter values and goodness of fit of the response-surface model describing the combined action of INH and RIF measured after 7 days**

| **Parameter** | **IMV** | **4MBE** |
| --- | --- | --- |
| E_max,INH_ (decline in log_10_ CFU/ml) | 5.11 [4.90; 5.33] | 4.41 [4.17; 4.67] |
| E_max,RIF_ (decline in log_10_ CFU/ml) | 6.33 [5.99; 6.67] | 5.91 [5.41; 6.41] |
| β_Emax_ (no unit) | -0.86 [-2.26; 0.54] | -3.73 [-6.76; -0.69] |
| H_INH_ (no unit) | 3.54 [2.64; 4.43] | 2.07 [1.74; 2.41] |
| H_RIF_ (no unit) | 0.79 [0.65; 0.93] | 1.12 [0.88; 1.35] |
| β_H_ (no unit) | 1.03 [-1.04; 3.09] | 1.67 [0.39; 2.95] |
| EC_50,INH_ (x MIC) | 0.53 [0.48; 0.58] | 0.91 [0.80; 1.03] |
| EC_50,RIF_ (x MIC) | 0.39 [0.30; 0.47] | 2.20 [1.61; 2.79] |
| Interaction parameter β_U50_ (no unit) | 1.43 [0.94; 1.91] | 0.68 [-0.38; 1.74] |
| **Goodness-of-fit** |  |  |
| Regression equation (obs. versus pred.) | y = 0.997x + 0.014 | Y = 0.99x + 0.045 |
| R^2^ | 0.90 | 0.96 |
| Mean prediction error (standard-deviation) | -0.0027 (0.63) | -0.016 (0.35) |
| Median absolute prediction error (%) | 12.2 | 12.5 |

Model parameters are described in the text Method section. obs., observed combined antibacterial effect; pred., model-based prediction of the combined. Parameter values are given as point estimate with [95% confidence interval]. MIC: minimum inhibitory concentration. RIF: Rifampicin; INH: Isoniazid
