## Supplementary material for "Rifampicin exposure reveals within-host *Mycobacterium tuberculosis* diversity in patients with delayed culture conversion": Table S3

**S3 Table. Primer sets for targeted NGS**

| **Primer name** | **Sequence** | **Size (bp)** |
| --- | --- | --- |
| 3280555-variant-F | GTGTAGCAGTAGCGGGCATT | 1695 |
| 3280555-variant-R | AATCGGGATTCTCGGTGAC |  |
| 3281868-variant-F | TGGACGACAGCATGAATAGC | 1748 |
| 3281868-variant-R | TGCAACGGATGTAGTGCTTC |  |
| 4145334-variant-F | CGGTTAAATGCTCCTCGAAA | 1785 |
| 4145334-variant-R | ACGGTGAATGGCAAGACTTC |  |
| 2579217-variant-F | CGCGATAGTCAAACAGCAAC | 1671 |
| 2579217-variant-R | CTGAGGGACAACGATGACAG |  |
| 649763-variant-F | CGGTCTACCAAACCGATGTT | 1647 |
| 649763-variant-R | TACAAGTCGGCCACTTTCCT |  |
| 3849212-variant-F | GAGCAACCAACTCTGGCTTC | 1921 |
| 3849212-variant-R | CCTGATCAAAGCTCGTCGTC |  |
