## Supplementary figures and images for "Rifampicin exposure reveals within-host *Mycobacterium tuberculosis* diversity in patients with delayed culture conversion"

### Figure S1

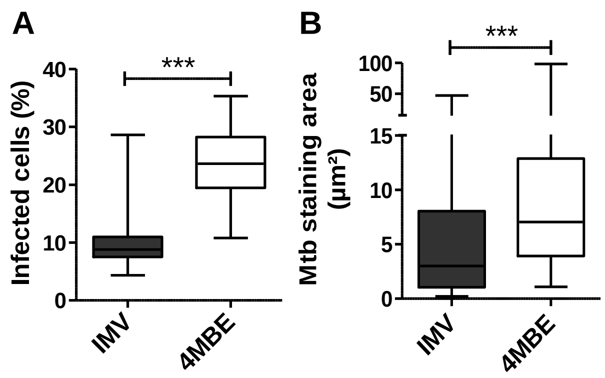

### Figure S2

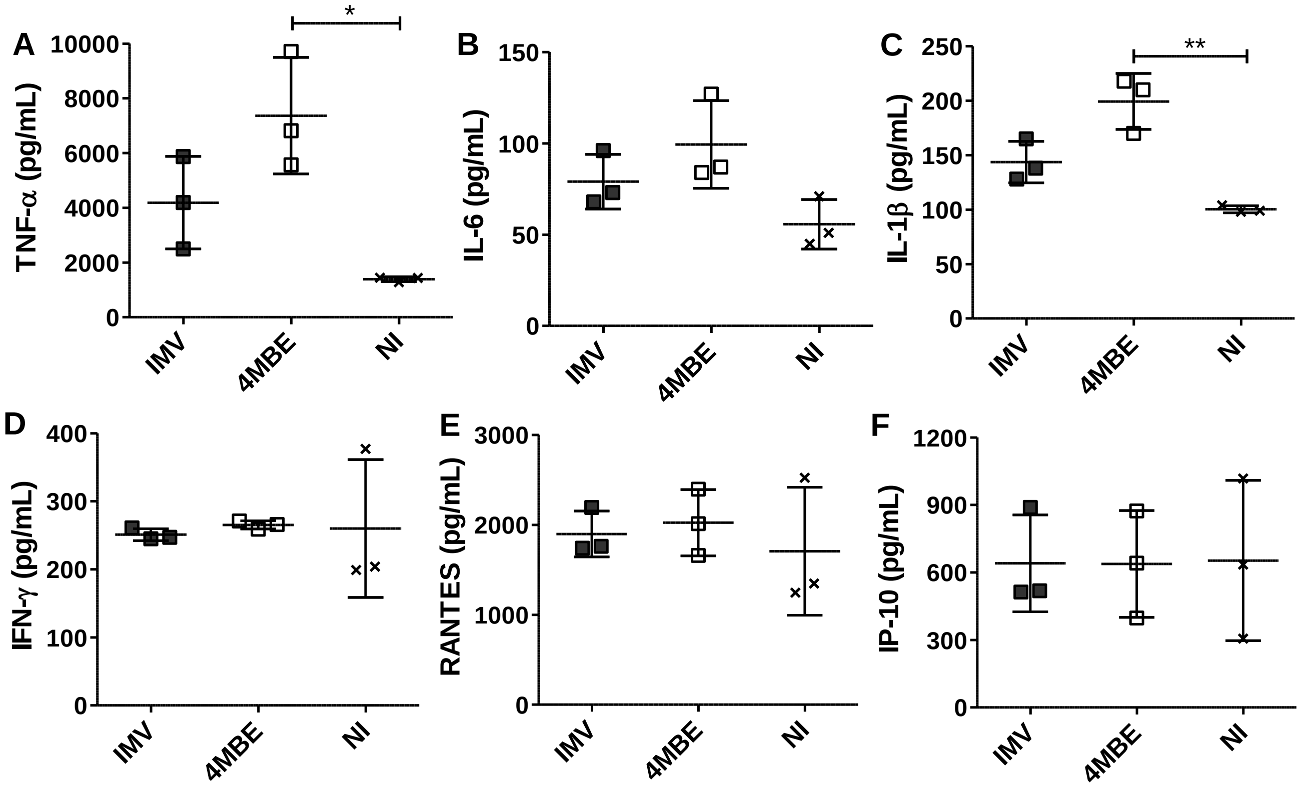

### Figure S3

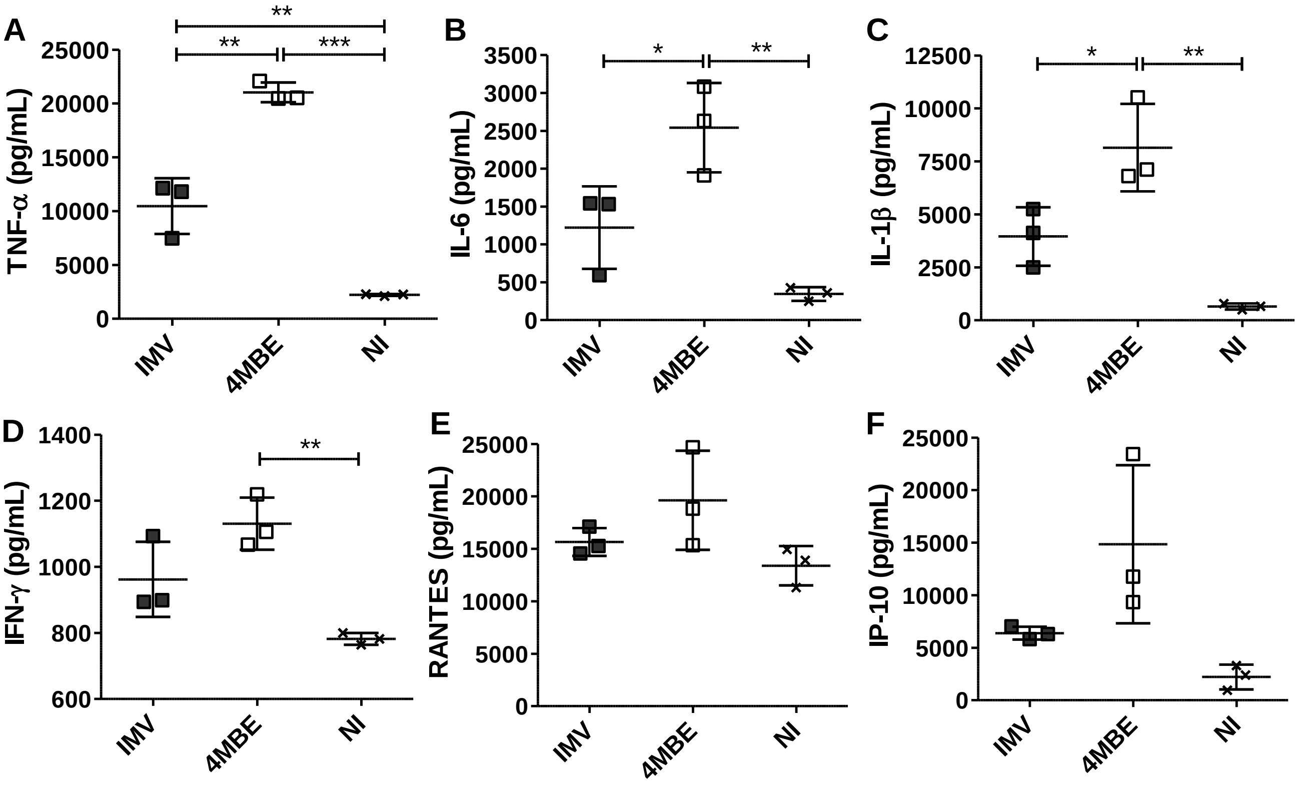
